# Reorganization of Human Brain Waves Across Diverse States of Consciousness

**DOI:** 10.64898/2026.05.27.728182

**Authors:** Panagiotis Fotiadis, Hyunwoo Jang, Rui Dai, Duan Li, Rodrigo Cofré, Christopher Timmermann, George A. Mashour, Anthony G. Hudetz, Zirui Huang

**Affiliations:** Department of Anesthesiology, University of Michigan Medical School, Ann Arbor, MI, USA; Center for Consciousness Science, University of Michigan, Ann Arbor, MI, USA; Neuroscience Graduate Program, University of Michigan, Ann Arbor, MI, USA; Michigan Psychedelic Center, University of Michigan Medical School, Ann Arbor, MI, USA; CRONOS Team, Inria Centre, Université Côte d’Azur, Sophia Antipolis, France; Centre for Consciousness Research, Department of Experimental Psychology, University College London, London, United Kingdom; Centre for Psychedelic Research, Department of Brain Sciences, Imperial College London, London, United Kingdom; Department of Pharmacology, University of Michigan Medical School, Ann Arbor, MI, USA

**Author notes:** Corresponding authors: Panagiotis Fotiadis, Zirui Huang.

## Abstract

Brain waves are ubiquitous phenomena of human brain activity. As they propagate, they coordinate neural communication, shaping conscious perception. Understanding how brain waves unfold across space and time is thus critical for uncovering the neural mechanisms that support and suppress consciousness. Here, we analyzed data from the Human Connectome Project alongside multiple independent human datasets of various states of consciousness collected during non-rapid eye movement sleep, propofol anesthesia, and psychedelic states produced by lysergic acid diethylamide, N,N-dimethyltryptamine, psilocybin, nitrous oxide, and ketamine. We then applied complex principal component analysis to map spatiotemporal propagation patterns of blood oxygen level-dependent activity across the human brain, under these diverse states of consciousness. We identified four dominant motifs of wave propagation: a global synchronized wave supporting unimodal-transmodal propagation, an anti-correlated unimodal-transmodal wave, an anti-correlated task-positive/task-negative wave, and an anti-correlated visual-somatomotor wave. Among them, the global wave exhibited the most pronounced state-dependent reconfiguration: in diminished states (sleep and anesthesia), the time needed for the wave to propagate across brain regions consistently increased and the distribution of regional contributions to the wave’s power became more spatially concentrated and heterogeneous across individuals, indicating slower, more fragmented, and less stereotyped dynamics. In contrast, propagation duration decreased under psychedelic states, reflecting accelerated global wave dynamics alongside a trend towards more spatially distributed and uniform regional contributions, consistent with a more integrated global wave propagation pattern. Beyond this global mode, diminished states slowed propagation primarily along the unimodal-transmodal axis, whereas psychedelic states selectively accelerated propagation along the task-positive/task-negative axis. Together, our findings reveal that diminished (sleep and anesthesia) and psychedelic states alter the spatiotemporal structure of wave propagation across the brain in opposite and distinct ways, providing a unifying account of how macroscale brain dynamics are dynamically reshaped under pharmacological and endogenous perturbations of consciousness.

## INTRODUCTION

Brain waves—spatiotemporal patterns of neural activity undulating across brain regions over multiple time scales—are increasingly recognized as key features enabling neural communication and coordination.^1,2^ These patterns can take multiple forms, ranging from standing wave-like activity in which a relatively fixed spatial pattern rises and falls in synchrony, to traveling waves in which peaks and troughs sequentially propagate across brain regions with systematic phase delays.^2^ Theoretical and experimental work suggests that wave dynamics can emerge naturally from recurrent neural circuits or be evoked by external stimuli^2^ and that as they travel, they transiently shape network excitability and spiking, coordinating distributed computation, and enabling flexible cognition.^3,4^ Consistent with this view, recent work has shown that unlike disorganized fluctuations in neural activity, spontaneous traveling waves in the visual cortex of awake primates can predict moment-to-moment fluctuations in perceptual detection, directly linking brain wave dynamics to conscious perception.^5^ A key question then emerges: if brain waves underlie neuronal coordination and conscious perception, how does their spatiotemporal organization change when levels of consciousness diminish, as in the case of sleep and anesthesia, or when conscious content is profoundly altered, as during exposure to psychedelic substances?

Growing electrophysiological work has found that conscious-unconscious transitions are accompanied by large-scale reconfigurations of wave dynamics. Transition to non-rapid eye movement sleep, for instance, is characterized by the emergence of reproducible slow oscillations (< 1 Hz) that traverse the cortex as anterior-to-posterior traveling waves, whose rate of occurrence increases as sleep deepens.^6^ Likewise, propofol anesthesia produces traveling waves resembling those observed during sleep;^7^ in addition to amplifying slow delta waves (∼ 1 Hz) and rendering them more spatially ordered, propofol also disrupts higher frequency traveling patterns (8 – 30 Hz) associated with conscious processing.^8,9^

Psychedelic states also profoundly alter brain wave dynamics. Electrophysiological studies show that serotonergic psychedelics such as N,N-dimethyltryptamine (DMT) and 5-methoxy-DMT reorganize cortical traveling waves. DMT induces propagation patterns of cortical activation resembling those elicited by visual stimulation, increasing posterior-to-anterior wave propagation while reducing anterior-to-posterior propagation typical of resting wakefulness.^10^ Instead, 5-MeO-DMT, while not producing consistent directional shifts in propagation, disorganizes spatiotemporal wave patterns rendering them unable to travel up and down the cortical hierarchy.^11^ The macroscale reorganization of traveling waves under the influence of other psychedelic states, however, remains largely unexplored.

Despite these advances in characterizing wave dynamics across both diminished and psychedelic states of consciousness, two important limitations remain. First, direct investigations of wave dynamics in diminished versus psychedelic states of consciousness are currently scarce. Second, most existing studies of wave propagation have predominantly relied on electrophysiology which, although temporally precise, provides limited spatial resolution compared to functional magnetic resonance imaging (fMRI). At the same time, the frequency- and region-dependent relationship between EEG-derived oscillations and fMRI-derived infra-slow fluctuations^12,13^ highlights that these imaging modalities capture complementary aspects of spatiotemporal dynamics. Therefore, extending temporally precise electrophysiological findings across multiple frequency bands with spatially resolved fMRI measurements of infra-slow activity could help paint a more complete picture of the full spatiotemporal landscape of brain wave dynamics across altered states of consciousness.

Here, we address these gaps in knowledge by combining whole-brain fMRI with a complex principal component analysis (CPCA) framework to map the brain’s spatiotemporal wave dynamics across diverse states of consciousness. Building on recent work showing that this approach can extract repeating, brain-wide propagation patterns (also referred to as waves) from fMRI signals,^14,15^ we analyzed data from the Human Connectome Project (HCP) alongside several independent datasets of altered states of consciousness ranging across wakefulness, diminished states (three propofol cohorts spanning graded depths and one sleep dataset capturing lighter and deeper stages), and psychedelic states (classical: lysergic acid diethylamide [LSD], DMT, psilocybin, and non-classical: nitrous oxide, ketamine). Using this methodology, we identified dominant waves of brain-wide signal propagation and quantified how their spatiotemporal properties change across states of consciousness, revealing global wave patterns that differentiate diminished from psychedelic states.

## RESULTS

### Dominant patterns of spatiotemporal propagation

To identify the changes in spatiotemporal propagation across different states of consciousness using multiple datasets, we first established a set of spatiotemporal reference patterns using the HCP dataset—a large, high-quality resting-state fMRI dataset commonly used as a normative reference for functional activity.^16^ The reference patterns were used to establish a common basis for the characterization of wave changes within a unified analytical framework. Specifically, we used CPCA to decompose the BOLD signal into a series of spatiotemporal modes, each characterized by a spatial pattern and its associated temporal dynamics (**Methods: Extraction of spatiotemporal propagation patterns**). We applied CPCA to the HCP data (*N* = 100 unrelated individuals) and retained the first four complex principal components as reference patterns (henceforth referred to as CPC1-CPC4), as they collectively accounted for approximately half (48%) of the total variance of the data (**Supplementary Figure 1**). This decision was further supported by the explained variance scree plot showing an elbow effect after the fourth complex component (**Supplementary Figure 1**), and by prior studies using a similar number of components.^14,15^

We next characterized the patterns’ propagation properties (**Figures 1**, **2**). To that end, brain regions were first grouped into seven canonical resting-state functional cortical networks (visual, somatomotor, dorsal attention, ventral attention, limbic, frontoparietal, and default mode) spanning the unimodal-transmodal hierarchy,^17,18^ as well as several subcortical networks (hippocampus, amygdala, basal ganglia, and thalamus) (**Methods: fMRI Processing**). The first complex principal component (CPC1) reflected a global activation/deactivation wave, wherein all brain networks were largely synchronized with one another (**Figure 1: CPC 1**). Consistent with this interpretation, removal of the global signal from the fMRI data eliminated this CPC1 wave (**Supplementary Analysis 1**). Besides temporally tracking the global signal (correlation between CPC1’s time course and the mean whole-brain signal over time: Pearson’s *r*=0.99), this wave’s spatial amplitude was driven by unimodal (i.e., visual and somatomotor) and attentional (i.e., dorsal and ventral attention) rather than transmodal (i.e., frontoparietal and default mode) and subcortical networks (permutation tests shown in **Figure 1: CPC 1** and **Supplementary Figure 2**). Computation of circular phase differences between each network’s phase and that of the whole brain (lead-lag analysis) revealed that CPC1, although largely synchronous, also expressed a structured propagation trajectory. Within this trajectory, activity progressed along a unimodal-transmodal axis, moving from somatomotor, to attentional and visual, then to default mode, and finally to frontoparietal networks (**Figure 2: CPC 1**). This observation was recapitulated by a strong correlation between CPC1 and a previously established unimodal-transmodal gradient (Gradient 1:^19^ Pearson’s *r*=0.82; **Supplementary Figure 3: Complex Principal Component 1**; **Methods: Correlation between spatiotemporal patterns and functional gradients**). Subcortical structures occupied distinct positions along this trajectory, with the amygdala leading the wave, and the basal ganglia and thalamus lagging towards the end of the wave cycle.

**Figure 1.**
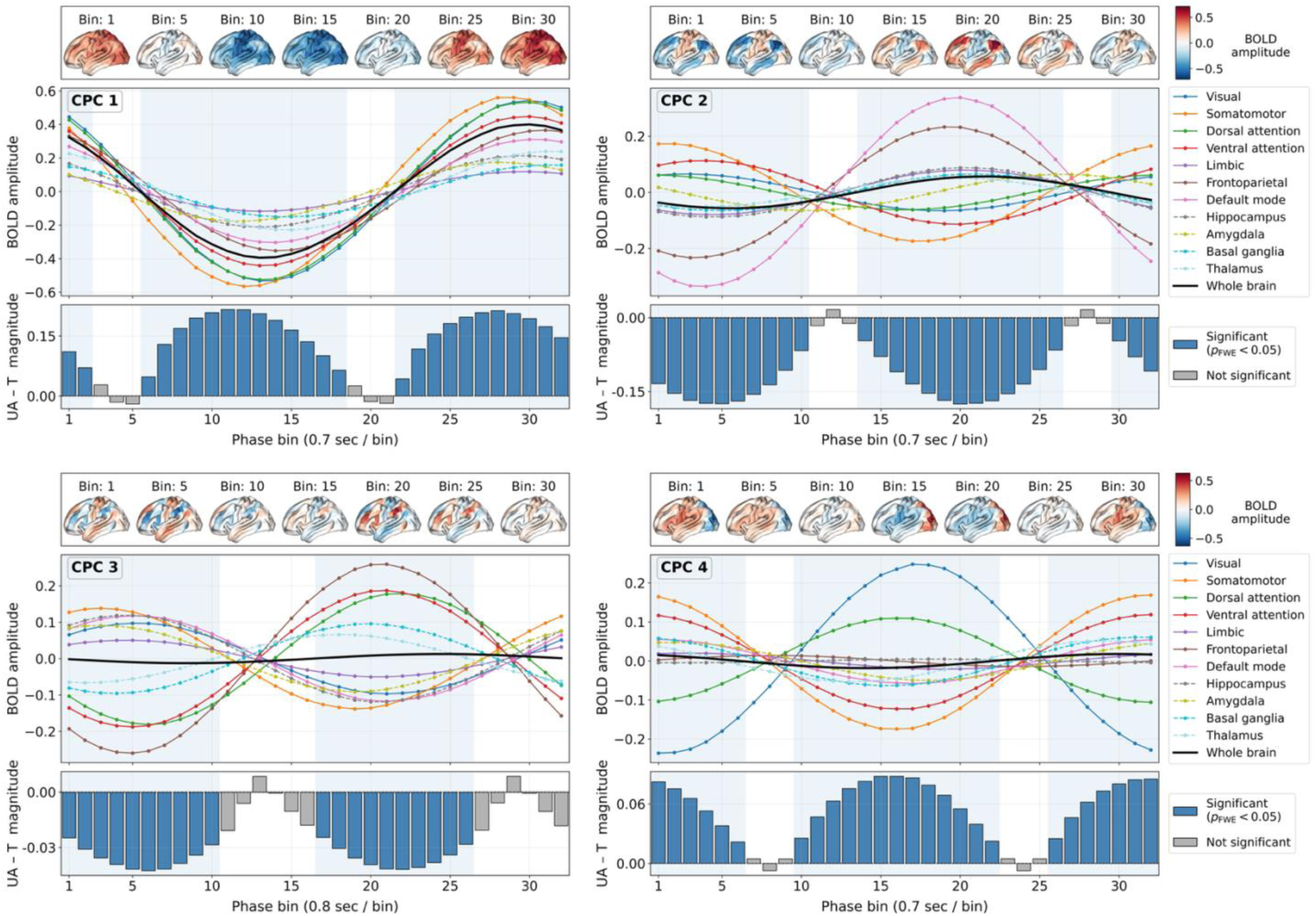
—Dominant spatiotemporal propagation patterns of brain activity. Each sub-figure corresponds to one of the four reference spatiotemporal propagation patterns of brain activity, derived from the HCP dataset. For each sub-figure, we display the pattern’s propagation motif at different points (i.e., phase bins) across the oscillatory cycle (**top**), each network’s phase-binned BOLD amplitude timecourses extracted from the complex principal component analysis reconstruction of the fMRI signal (**middle**), and the unimodal and attentional versus transmodal magnitude dominance at each phase bin, computed as the difference between the mean absolute reconstructed BOLD amplitude across unimodal/attentional (UA: visual, somatomotor, dorsal attention, and ventral attention) and transmodal (T: frontoparietal and default mode) regions, at each phase bin (**bottom**). Statistical significance at each phase bin was determined via permutation testing where the unimodal/attentional and transmodal labels were randomly shuffled across brain regions to generate a null distribution of unimodal/attentional and transmodal differences; the empirical difference at each bin was then compared against this null distribution, and *p*-values were corrected for multiple comparisons across bins (Family-Wise Error; *p_FWE_*). Each phase bin’s duration in seconds was also approximated and provided in the x-axis label. CPC: Complex Principal Component.

**Figure 2.**
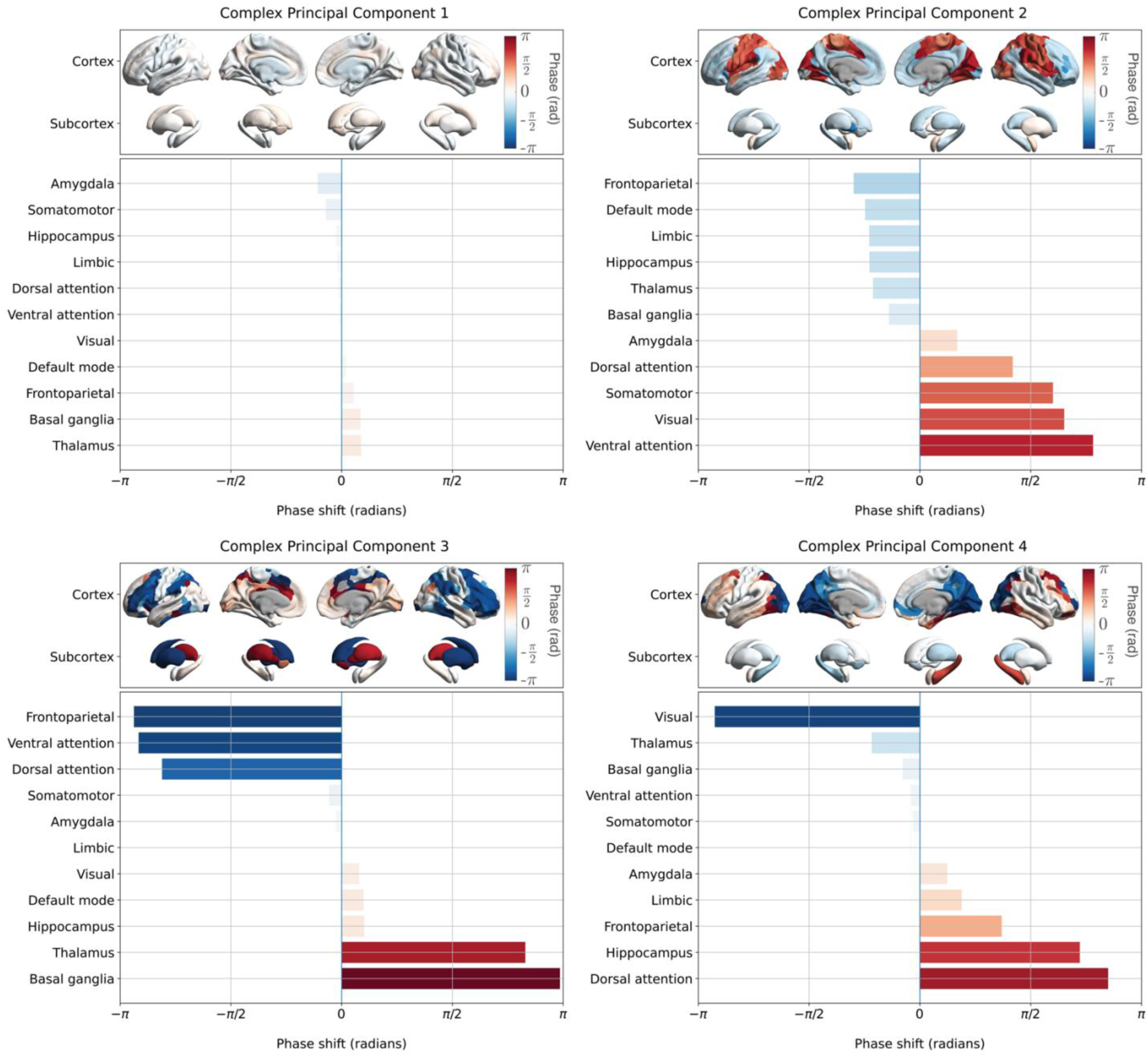
—Directional organization of spatiotemporal propagation patterns. Each sub-figure corresponds to one of the four reference spatiotemporal propagation patterns of brain activity, derived from the HCP dataset. For each sub-figure, we display the pattern’s directional organization as derived from phase lead-lag analyses: for each complex principal component, each brain region’s phase offset was computed relative to the whole brain as the circular mean difference between the region’s mean phase and that of the whole brain’s. Using this information, we generated ‘phase delay maps,’ where each brain region was assigned its corresponding phase shift value: colder colors indicate brain regions leading the cycle relative to the global phase, while warmer colors indicate brain regions lagging behind the global phase (**top** within each subplot). We next computed the average phase shifts across networks: horizontal bars show different networks’ corresponding phase shift (radians) within an oscillatory cycle within the [−*π*, *π*) range, ordered such that networks with negative phase shifts are leading the propagation pattern, while networks with positive phase shifts are lagging behind the global phase (**bottom** within each subplot).

Interestingly, the second complex principal component (CPC2) revealed a different type of unimodal-transmodal propagation motif, characterized by an anti-correlated, anti-phasic organization (**Figures 1, 2: CPC 2**). Within this pattern, unimodal and transmodal cortices not only fluctuated out of phase, but occupied opposing extrema of the propagation cycle, such that periods of maximal activity in unimodal regions coincided with troughs in activity across transmodal regions, and *vice versa*, forming a reciprocal wave alternating between the two systems. Similar to CPC1, CPC2 also exhibited a strong spatial correlation with Gradient 1 (Pearson’s *r*=0.9; **Supplementary Figure 3: Complex Principal Component 2**). In contrast to CPC1, however, the spatial amplitude of this unimodal-transmodal pattern was driven by transmodal rather than unimodal or attentional networks (permutation tests shown in **Figure 1: CPC 2** and **Supplementary Figure 2**).

Next, the third complex principal component (CPC3) captured a spatiotemporal pattern propagating between task-positive (i.e., frontoparietal and attentional) and task-negative (i.e., default mode network) networks (**Figures 1, 2: CPC 3**), a result validated by a strong correlation between CPC3 and a previously established task-positive/task-negative gradient (Gradient 3;^19^ Pearson’s *r*=0.91; **Supplementary Figure 3: Complex Principal Component 3**). Within this pattern, subcortical structures such as the thalamus and the basal ganglia were relatively in-phase with the task-positive end of this axis (**Figures 2: CPC 3**). Similar to CPC2, this motif exhibited an anti-correlated, anti-phasic organization between the two opposing ends of the propagation trajectory. Lastly, the fourth complex principal component (CPC4) reflected a wave propagating between visual and somatomotor (as well as default mode) networks in an anti-correlated, anti-phasic pattern (**Figure 1, 2: CPC 4**). As with the previous CPCs, there was a strong correlation between this CPC and a previously established visual-somatomotor gradient (Gradient 2;^19^ Pearson’s *r*=0.86; **Supplementary Figure 3: Complex Principal Component 4**).

### Properties of spatiotemporal propagation

Afterwards, we analyzed fMRI data collected from datasets spanning wakefulness, diminished states (non-rapid eye movement [NREM] sleep, propofol anesthesia), as well as psychedelic states of consciousness (LSD, DMT, psilocybin, ketamine, and nitrous oxide). We specifically applied the CPCA framework to each subject’s time-series, separately for each state of consciousness and dataset. To directly compare spatiotemporal propagation properties across datasets, we identified the spatiotemporal patterns of each subject, state, and dataset that best matched the first four HCP-derived reference patterns using a circular correlation approach, and subsequently aligned them to their reference (**Methods: Extraction of spatiotemporal propagation patterns**). The group-level alignment yielded high similarity between our aligned spatiotemporal patterns and the HCP-derived references (**Supplementary Figure 4**), highlighting the method’s reliability and generalizability. This process resulted in a comparable set of spatiotemporal patterns for each subject and state of consciousness (as statistically validated in **Supplementary Figure 4**).

We next compared the patterns’ spatiotemporal characteristics, focusing specifically on (i) the traveling component duration of each spatiotemporal pattern, that is, the portion of the pattern’s overall duration attributed to spatial phase propagation, and (ii) how concentrated each spatiotemporal pattern’s spatial power is across brain regions.

#### A. Diminished states of consciousness exhibit prolonged traveling component durations

Each spatiotemporal pattern’s overall duration was first estimated in seconds (**Figure 3**; **Methods: Spatiotemporal patterns’ overall duration**). We then computed the pattern’s traveling component duration, that is the time required for the wave to traverse the range of phase offsets observed across brain regions, as different regions reach their fMRI-derived blood oxygen level-dependent (BOLD) signal peaks at different time-points. To do so, we computed the range of spatial phase angles that brain regions occupied within that wave and multiplied that range by the wave’s overall duration. Thus, for instance, if a given CPC captured a pure traveling wave, its spatial phase range would span the whole range of available phases (0 to 2*π*), yielding a traveling component duration equal to the overall duration of the wave; in contrast, if the CPC captured a predominantly standing wave, its spatial phase range would be extremely narrow, leading to a traveling component duration close to zero.

**Figure 3.**
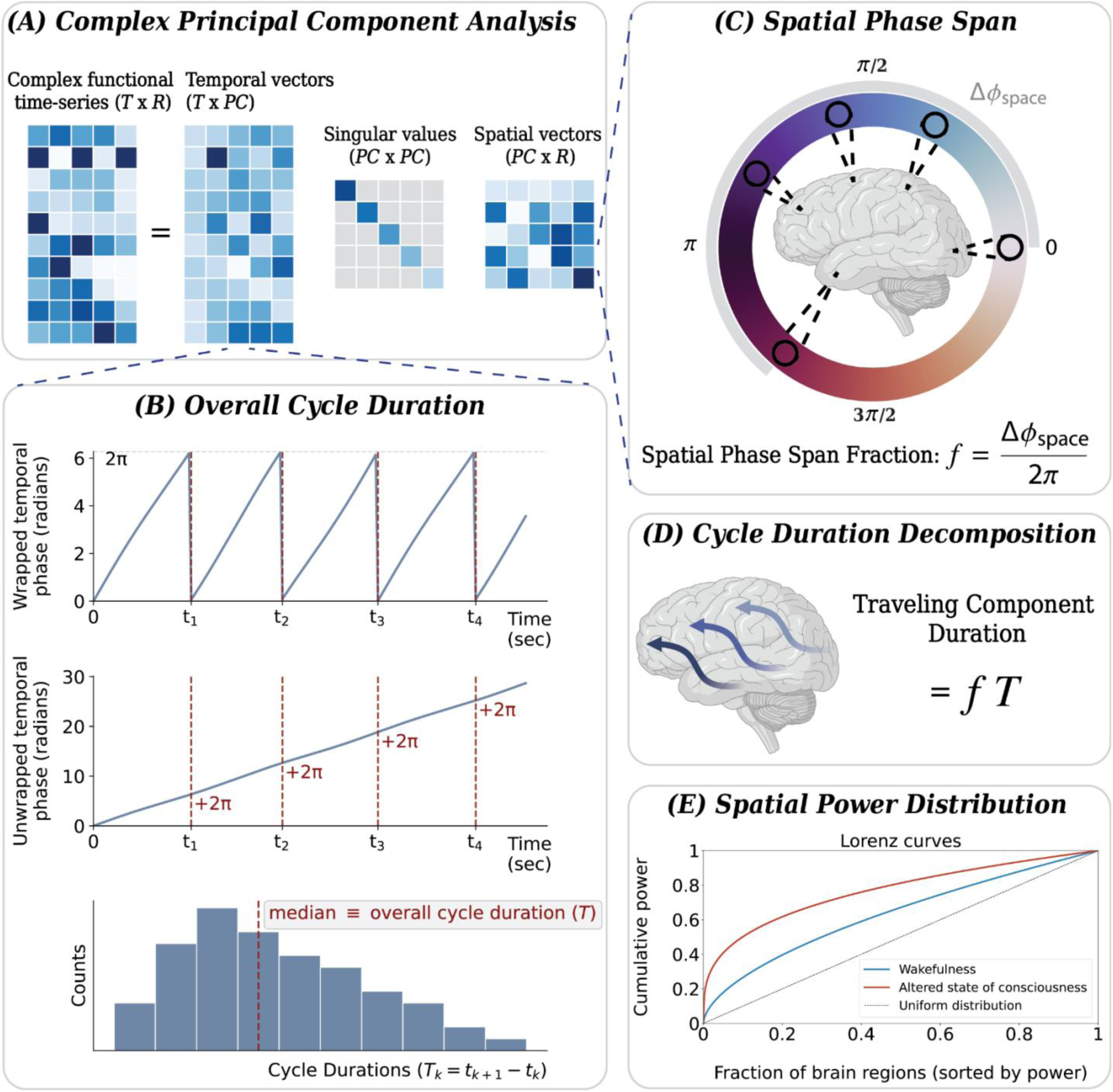
—Computation of spatiotemporal patterns’ traveling component duration and spatial power distribution. (A) The fMRI signal time-series were first Hilbert-transformed into their analytic complex version, and decomposed into paired temporal and spatial components using complex principal component analysis. (**B**) To derive each complex principal component’s temporal duration, the instantaneous phase (radians) of its corresponding temporal vector was extracted and temporally unwrapped to yield a continuous phase trajectory; a full propagation cycle was defined every time an integer multiple of 2*π* was crossed. Cycle durations were then computed as the temporal differences between consecutive crossings; the median of these values was defined as the overall cycle duration of the spatiotemporal pattern and multiplied by the fMRI scan’s repetition time to convert the value from radians to seconds. (**C**) Each principal component’s spatial phase range was computed from its corresponding spatial vector by estimating the median phase span across brain regions (re-referenced: see **Methods**), expressed as a fraction of a full oscillatory cycle. (**D**) The traveling component duration of the propagation pattern was then defined as its spatial phase fraction multiplied by its overall duration. (**E**) The Lorenz curve for each state represents the cumulative fraction of the total power (y-axis) of a spatiotemporal pattern captured by the top fraction of brain regions (x-axis), after sorting regions within each state by descending power contribution. The dashed diagonal denotes a uniform distribution wherein all regions contribute equally to the spatiotemporal pattern’s overall power. Curves deviating further away from the diagonal indicate greater concentration of power within a smaller subset of regions. The area under the curve for each state is quantified as the Gini coefficient (see **Methods**).

To assess how the traveling component duration of each spatiotemporal pattern (CPC1-CPC4) changed from wakefulness to diminished states of consciousness, we leveraged four independent datasets capturing reduced levels of consciousness, induced either endogenously via sleep, or pharmacologically via propofol anesthesia. The sleep dataset consisted of individuals who had fMRI data acquired during wakefulness and during transitions into light (N1) and deeper (N2) sleep stages. In parallel, the propofol datasets spanned graded levels of anesthesia, ranging from lighter sedation (Propofol Dataset 1: 1 μg/mL effect-site concentration [PPF: 1.0]), to intermediate sedation (Propofol Dataset 1: 1.9 μg/mL [PPF: 1.9] and Propofol Dataset 2: 2.2 μg/mL [PPF: 2.2]), and finally to deeper sedation (Propofol Dataset 3: 2.7 μg/mL [PPF: 2.7]).

We began by quantifying effect size differences (paired Hedge’s *g*; **Methods: Statistical Analyses**) in traveling component duration across the four spatiotemporal patterns, between each altered state and its corresponding baseline wakefulness. All diminished states were notably associated with a robust increase in CPC1 traveling component duration (**Figure 4**). The magnitude of this effect largely scaled with depth of sleep and anesthesia, ranging from light sleep (N1: *g*=0.9; CI=[0.4, 1.4]; *p_FDR_*=0.020) and light propofol sedation (PPF 1.0: *g*=0.8; CI=[0.1, 1.3]; *p_FDR_*=0.049), to deeper sleep (N2: *g*=1.2; CI=[0.5, 2.1]; *p_FDR_*=0.020) and intermediate propofol sedation (PPF 1.9: *g*=1.4; CI=[0.7, 2.6]; *p_FDR_*=0.012, PPF 2.2: *g*=0.9; CI=[0.3, 1.5]; *p_FDR_*=0.020), reaching its largest increase under deep propofol sedation (PPF 2.7: *g*=2.4; CI=[1.5, 3.3]; *p_FDR_*<10^-5^). The traveling component durations of the remaining spatiotemporal patterns CPC2-CPC4 also tended to increase during diminished states of consciousness, although less strongly than CPC1. Paired within-subject comparisons corroborated these results, reporting significant increases in traveling component durations across all CPCs (**Figure 5A, B**), with the largest effect observed once again in CPC1 (*t*=8.8; *p_holm_*=1.9×10^-12^).

**Figure 4.**
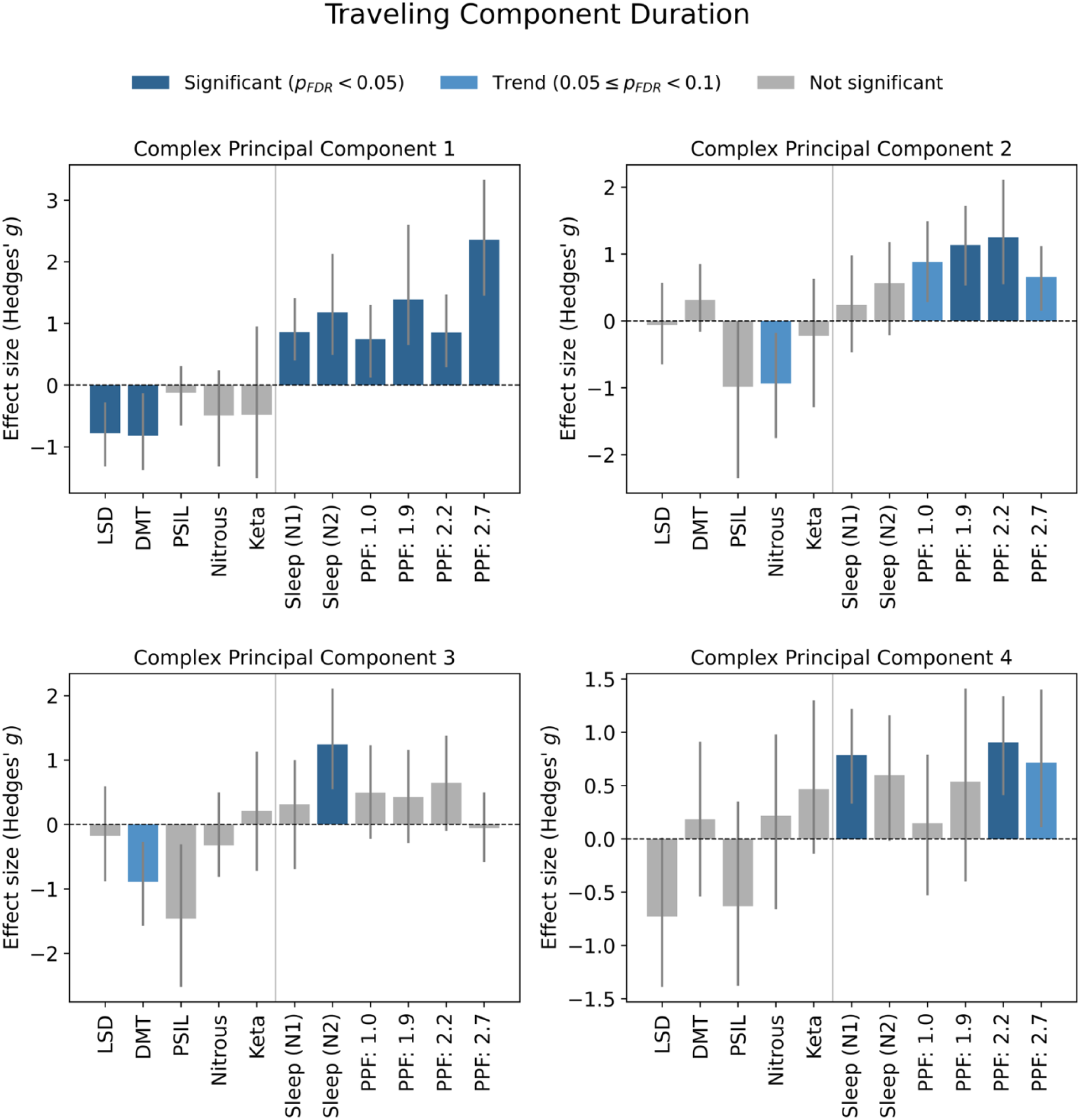
—Effect size differences in traveling component duration for each spatiotemporal pattern. For each complex principal component, we show the within-subject difference in traveling component duration (in seconds) between each altered-state-of-consciousness dataset and its respective wakefulness state. Vertical bars were computed using paired standardized mean differences (Hedge’s *g*), along with bootstrapped (95%) confidence intervals (grey lines). Significance was assessed using paired *t*-tests, corrected for multiple comparisons across datasets (Benjamini-Hochberg false-discovery rate; *p_FDR_*) and color-coded based on significance levels (see **legend**).

**Figure 5.**
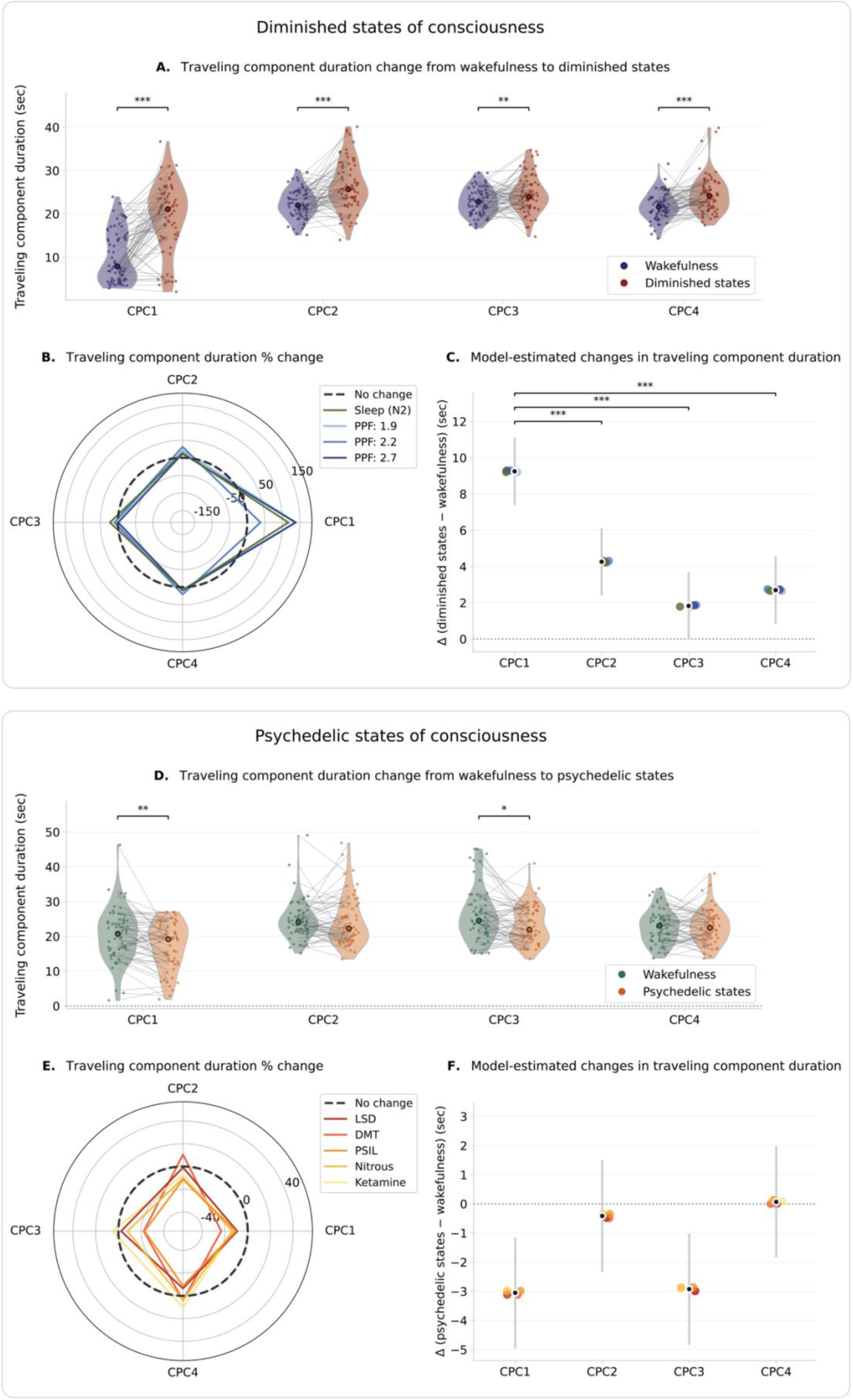
—Within-subject and mixed-effects estimates of traveling component duration. **(A, D):** Violin plots show within-subject changes in traveling component duration from wakefulness to the altered state (A: diminished; D: psychedelic), for each complex principal component (CPC1-CPC4). Paired individual observations aggregated across all included datasets are displayed as datapoints (A: 4 datasets; 72 total participants; D: 5 datasets; 61 participants) and connected for each individual with a line; median values are also indicated as thicker datapoints. Statistical significance was assessed using paired *t*-tests across subjects for each CPC and corrected for multiple comparisons (Holm; *p_holm_*): #: 0.05 ≤ *p_holm_* < 0.10; *: *p_holm_* < 0.05; **: 0.001 ≤ *p_holm_* < 0.01; ***: *p_holm_* < 0.001. **(B, E):** Spider plots (or radar plots) showing the percent change in across-subjects mean traveling component duration from wakefulness to each altered state, across each CPC. **(C, F):** Forest plots summarizing the mixed-effects modeling estimates of traveling component duration change from wakefulness to altered states, for each CPC. The black datapoints and grey vertical bars respectively denote the fixed-effect estimates of the model and their corresponding 95% confidence intervals; the colored datapoints represent each dataset’s random time slope (color matches the legend of sub-figures B and E, respectively). Horizontal lines and asterisks indicate when the model-estimated change in CPC1 was significantly larger than that of the other CPCs, corrected for multiple comparisons (Holm; *p_holm_*). *: 0.01 ≤ *p_holm_* < 0.05; **: 0.001 ≤ *p_holm_* < 0.01; ***: *p_holm_* < 0.001.

To formally test whether these effects in traveling component duration were consistent across sleep and propofol anesthesia, we fit a linear mixed-effects model in which state (pre: wakefulness vs post: diminished state), principal component classification (CPC1 through CPC4), and their interaction were included as fixed effects; dataset-specific random intercepts and random slopes for each state were also specified to account for variability across datasets (4 datasets; 72 total participants; 576 observations; **Methods: Statistical Analyses**). Model-estimated wakefulness-diminished state contrasts confirmed that traveling component duration increased overall, though to different extents, across spatiotemporal patterns: CPC1 showed the largest increase (*β*=9.3 sec; CI=[7.4, 11.1]; *p_holm_*<10^-5^), followed by CPC2 (*β*=4.3 sec; CI=[2.4, 6.1]; *p_holm_*=2.2×10^-5^) and CPC4 (*β*=2.7 sec; CI=[0.8, 4.6]; *p_holm_*=0.009), whereas CPC3 showed a borderline, albeit non-significant, increase (*β*=1.8 sec; CI=[-0.04, 3.7]; *p_holm_*=0.055) (**Figure 5C**). Interaction contrasts further confirmed that the pre-post increase was significantly larger for CPC1 compared to CPC2-CPC4 (state x CPC interaction: all *p*<10^-5^; reference: CPC1), a result mirrored by pairwise contrasts between slopes (**Figure 5C**: CPC1 > CPC2-CPC4: all *p_holm_*≤5.9×10^-5^), indicating that propagation slowing was dominated by CPC1. Importantly, dataset-specific slopes clustered around the fixed-effect estimates for each CPC with minimal between-dataset variability in time slopes (variance=0.006; **Figure 5C**), suggesting that all datasets displayed very similar pre-post increases in traveling component duration.

Given that traveling component duration was defined as the product of overall cycle duration and spatial phase range, we next determined which factor contributed to the observed increases in CPC1 traveling component duration by examining each constituent separately using mixed-effects modeling. Both measures showed significant increases during diminished states of consciousness: the overall cycle duration of CPC1 increased (*β*=3.7 sec; CI=[1.9, 5.5]; *p_holm_*=1.3×10^-4^; **Supplementary Figure 5A-C)**, as did the spatial phase range spanned by brain regions (*β*=0.3; CI=[0.2, 0.4]; *p_holm_*=9.8×10^-10^; **Supplementary Figure 6A-C**). Therefore, the observed lengthening of CPC1 traveling component duration likely reflected a synergistic effect between slower oscillatory cycles and a broader spatial phase dispersion across brain regions.

Collectively, reduced levels of consciousness were characterized by robust increases in traveling component duration dominated by the global pattern CPC1 and followed by the unimodal-transmodal pattern CPC2, with no reliable change in task-positive/task-negative pattern CPC3’s traveling component duration.

#### B. Psychedelic states of consciousness exhibit shorter traveling component durations

In contrast to diminished states of consciousness, psychedelic states displayed reductions in CPC1 traveling component duration, particularly under the classical psychedelics LSD (*g*=-0.8; CI=[-1.3, −0.3]; *p_FDR_*=0.037) and DMT (*g*=-0.8; CI=[-1.4, −0.1]; *p_FDR_*=0.037); duration changes in the remaining spatiotemporal patterns were less consistent (**Figure 4**). Paired within-subject comparisons complemented these results, also revealing significant reductions in CPC1 (*t*=-3.5; *p_holm_*=0.004) and CPC3 (*t*=-3.0; *p_holm_*=0.012) traveling component durations (**Figure 5D, E**).

To determine whether these effects were consistent across psychedelic datasets, we once again used linear mixed-effects modeling (5 datasets; 61 participants; 488 observations). Consistent with the effect size and paired comparisons, model-estimated pre-post contrasts confirmed significant reductions in traveling component duration for CPC1 (*β*=-3.1 sec; CI=[-5.0, −1.1]; *p_holm_*=0.007) and CPC3 (*β*=-2.9 sec; CI=[-4.8, −1.0]; *p_holm_*=0.008) (**Figure 5F**). To further characterize these changes, we separately examined the two constituents of traveling component duration: overall cycle duration and spatial phase range. Decreases in CPC1 traveling component duration were primarily driven by a narrowing of the spatial phase range across psychedelic states (*β*=-0.07; CI=[-0.1, −0.04]; *p_holm_*=4.2×10^-4^; **Supplementary Figure 6D-F**), whereas decreases in CPC3 traveling component duration were primarily driven by the shortening of the wave’s overall cycle duration (*β*=-2.9 sec; CI=[-4.8, −1.1]; *p_holm_*=0.008; **Supplementary Figure 5D-F)**. As with the diminished datasets, dataset-specific time slopes clustered closely around the fixed-effect estimates (variance=0.004; **Figure 5F**), indicating consistent decreases in traveling component duration across the examined psychedelic compounds.

Together, these results demonstrate that psychedelic states selectively shorten the traveling component durations of both global pattern CPC1 and task-positive/task-negative pattern CPC3—accelerating the wave dynamics along these axes—without impacting the unimodal-transmodal pattern CPC2.

#### C. Diminished states of consciousness increase spatial concentration of traveling wave power while promoting heterogeneous regional dominance across subjects

We next examined the extent to which each brain region contributed to the different propagation patterns’ spatial power, and how that power distribution differed between diminished and psychedelic states of consciousness. For this purpose, we computed the relative power that each brain region contributed to the overall spatiotemporal propagation’s power and estimated the Gini coefficient of that normalized regional power distribution at each state of consciousness (**Figure 3E**; **Methods: Spatial distribution of regional power**). Intuitively, lower Gini coefficients indicate that power is evenly distributed across brain regions, while higher coefficients indicate greater concentration of power within a smaller subset of regions.

We began by examining changes in Gini coefficients during the transition from wakefulness to diminished states of consciousness. Dataset-level effect size (paired Hedge’s *g*) analyses revealed a marked increase in CPC1 Gini coefficients during diminished states, suggesting that traveling wave power became more spatially concentrated (**Supplementary Figure 7**). As with CPC1 traveling component duration, the magnitude of change in CPC1’s Gini coefficient largely scaled with depth of unconsciousness, displaying moderate increases under deep sleep (N2: *g*=1.0; CI=[0.1, 1.8]; *p_FDR_*=0.097) and intermediate propofol sedation (PPF 1.9: *g*=1.0; CI=[0.4, 1.8]; *p_FDR_*=0.059, PPF 2.2: *g*=1.1; CI=[0.7, 1.8]; *p_FDR_*=3×10^-4^), and more pronounced increases under deeper propofol anesthesia (PPF 2.7: *g*=1.9; CI=[1.2, 2.5]; *p_FDR_*<10^-5^).

These effect-size results were also reflected by the regional power histogram distributions in CPC1 across diminished states. The subject-level regional power histograms revealed increasingly skewed redistributions of power in CPC1 with a smaller fraction of regions contributing a larger share of the total power of the wave, particularly across intermediate and deeper levels of propofol sedation (**Supplementary Figure 8**). To further qualify this redistribution, we plotted the Lorenz curves for wakefulness and diminished states and examined their within-subject differences (**Figure 6A**). Conceptually, the Lorenz curve describes how cumulative power is distributed across regions ranked by contribution, with greater curvature reflecting more unequal distributions. Accordingly, positive deviations in the Lorenz-difference curves indicate that a smaller fraction of regions accounts for a larger fraction of the total power. Consistent with the Gini coefficient results, decreasing levels of consciousness broadly exhibited larger and more extended deviations from zero in the CPC1 Lorenz-difference curves, as evinced by a progressively greater area under the curves (**Figure 6A**).

**Figure 6.**
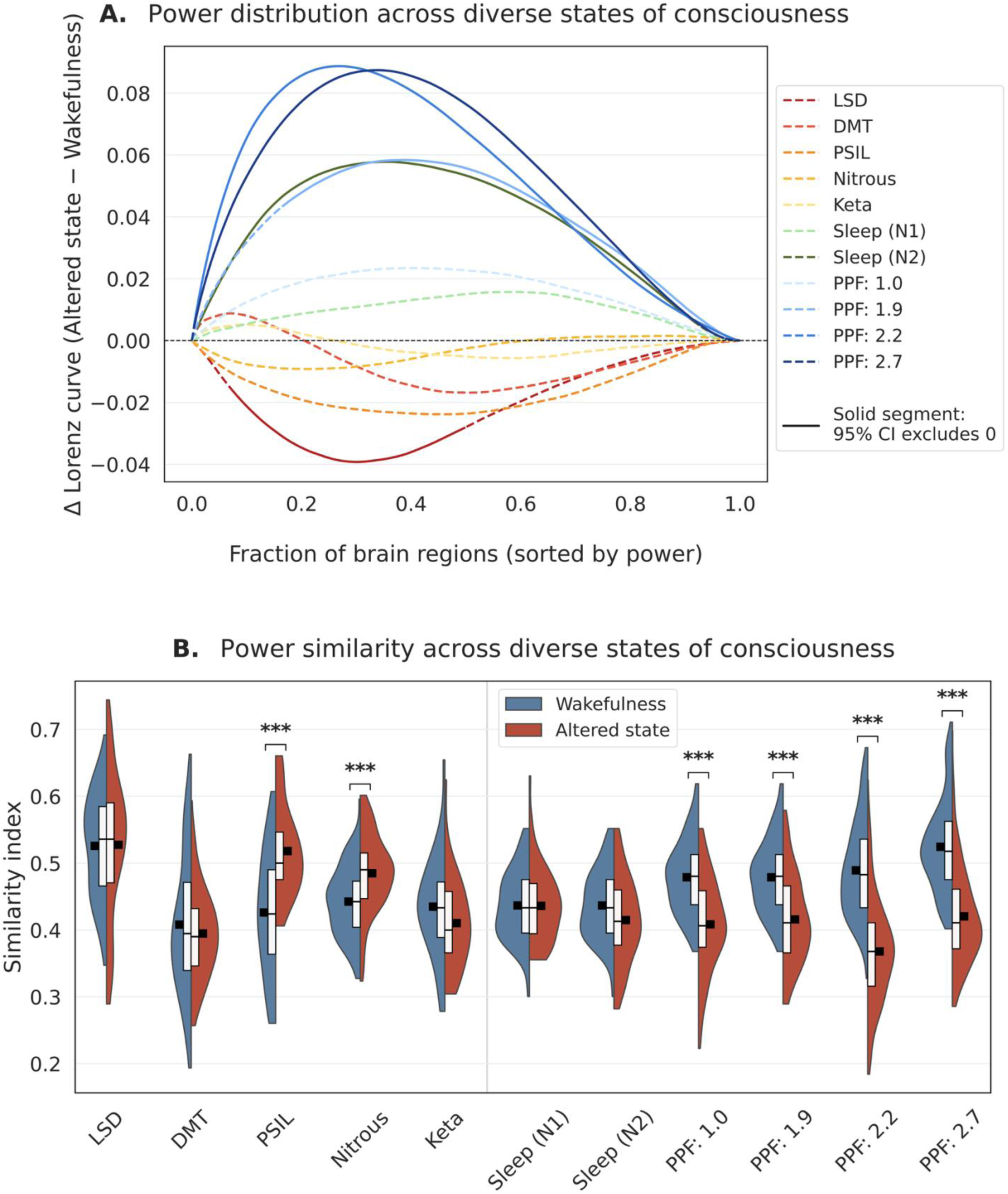
—Power distributions and similarity across states of consciousness. **(A) Power distribution—**For each dataset, we first generated the subject-level Lorenz curves of the normalized contribution of each brain region towards the overall CPC1 power, during wakefulness and during each altered state of consciousness. We then computed the difference (Lorenz-difference curves) between the two states for each subject, and plotted the average difference for each dataset. To identify the fraction of brain regions over which the spatial accumulation of power significantly differed across states, 95% confidence intervals were computed based on the *t*-distribution of the Lorenz-difference curves; curve segments where the confidence interval did not include zero were labeled as significant (solid line, instead of dashed line). **(B) Similarity in power distribution—**For each dataset, we showcase the distribution of pairwise Jaccard similarity values (Similarity index) quantifying the overlap in dominant brain regions across subjects during wakefulness (blue) and each altered state of consciousness (red). For each subject, dominant regions were defined as the ones with the top 50% of regional power contributions to the CPC1 wave. Pairwise similarity was then computed between all subject pairs within each state of consciousness (see **Methods: Similarity in spatial distribution of regional power**), with lower similarity indices indicating greater heterogeneity in the spatial distribution of dominant brain regions across subjects. Half violin plots show the distribution of similarity indices across all pairs of subjects within each state, white boxplots indicate the median and inter-quantile range, and the black squares indicate the mean similarity index for that state. Asterisks signify when the altered state demonstrated a similarity index significantly different from that of wakefulness, corrected for multiple comparisons (Benjamini-Hochberg false-discovery rate; *p_FDR_*). *: 0.01 ≤ *p_FDR_* < 0.05; **: 0.001 ≤ *p_FDR_* < 0.01; ***: *p_FDR_* < 0.001.

Paired within-subject comparisons expanded upon these findings, revealing significant increases in Gini coefficients predominantly in CPC1 (*t*=8.5; *p_holm_*=7.3×10^-12^), followed by CPC3 (*t*=3.1; *p_holm_*=0.007) and CPC4 (*t*=2.9; *p_holm_*=0.011) (**Figure 7A, B**). Mixed-effects model-estimated contrasts once again revealed a significant increase in CPC1 spatial inequality (*β*=0.1; CI=[0.05, 0.14]; *p_holm_*=6×10^-4^) during diminished states, with changes in CPC2-CPC4 not remaining significant (**Figure 7C**). Interaction contrasts further supported how the spatial redistribution observed in CPC1 was much more pronounced compared to all other CPCs (state × CPC: all *p*<10^-5^; reference: CPC1 and pairwise slope contrasts: CPC1 > CPC2-CPC4: all *p_holm_*≤9.4×10^-6^; **Figure 7C**). As before, dataset-specific slopes clustered tightly around the fixed-effect estimates with minimal between-dataset variability (variance=0.002; **Figure 7C**), indicating consistent increases in CPC1 spatial inequality across datasets.

**Figure 7.**
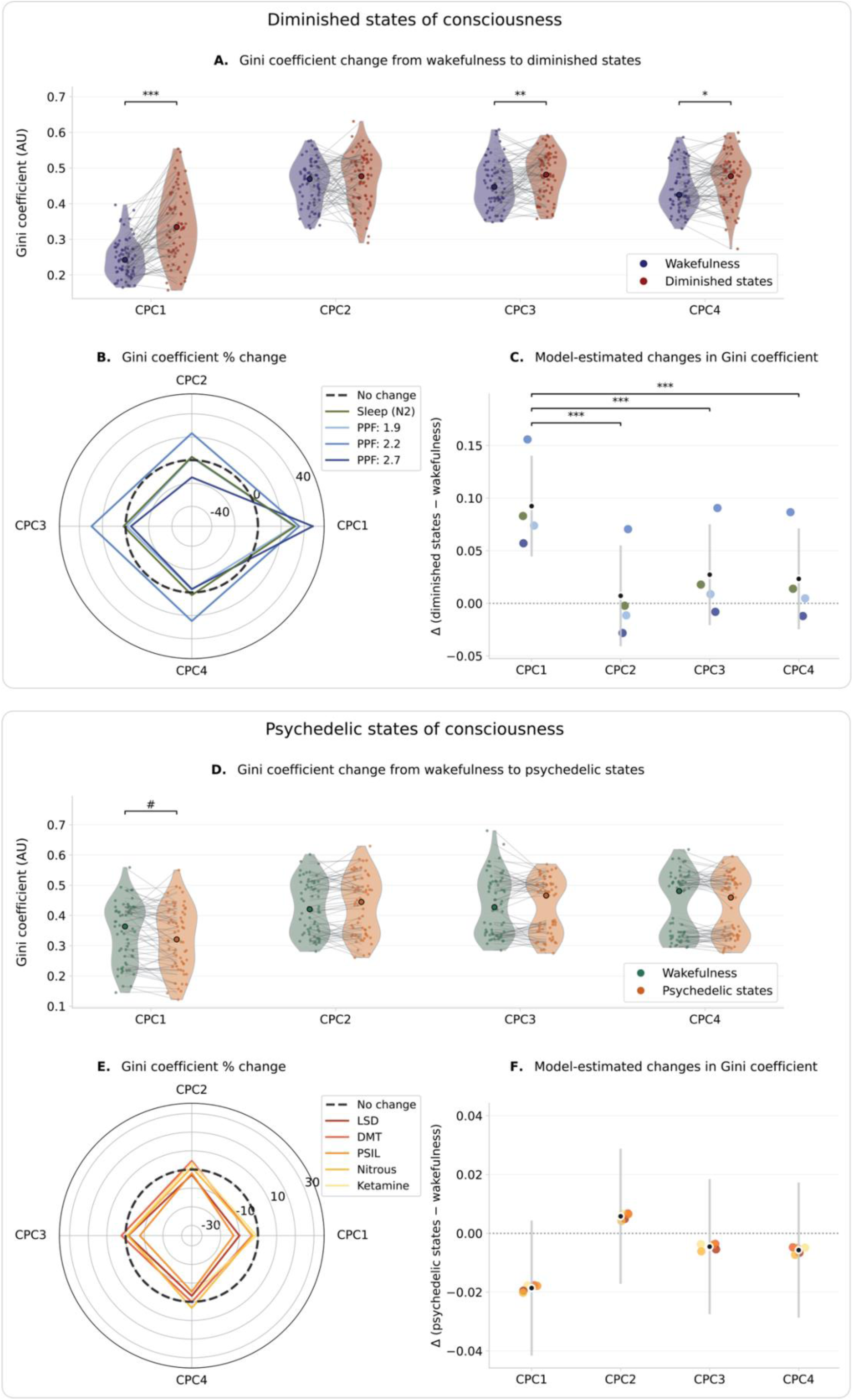
—Within-subject and mixed-effects estimates of Gini coefficient. **(A, D):** Violin plots show within-subject changes in the Gini coefficient from wakefulness to the altered state (A: diminished; D: psychedelic), for each complex principal component (CPC1-CPC4). Paired individual observations aggregated across all included datasets are displayed as datapoints (A: 4 datasets; 72 total participants; D: 5 datasets; 61 participants) and connected for each individual with a line; median values are also indicated as thicker datapoints. Statistical significance was assessed using paired *t*-tests across subjects for each CPC and corrected for multiple comparisons (Holm; *p_holm_*): #: 0.05 ≤ *p_holm_* < 0.10; *: 0.01 ≤ *p_holm_* < 0.05; **: 0.001 ≤ *p_holm_* < 0.01; ***: *p_holm_* < 0.001. **(B, E):** Spider plots (or radar plots) showing the percent change in the across-subjects mean Gini coefficient from wakefulness to each altered state, across each CPC. **(C, F):** Forest plots summarizing the mixed-effects modeling estimates of Gini coefficient change from wakefulness to altered states, for each CPC. The black datapoints and grey vertical bars respectively denote the fixed-effect estimates of the model and their corresponding 95% confidence intervals; the colored datapoints represent each dataset’s random time slope (color matches the legend of sub-figures B and E, respectively). Horizontal lines and asterisks indicate when the model-estimated change in CPC1 was significantly larger than that of the other CPCs, corrected for multiple comparisons (Holm; *p_holm_*). *: 0.01 ≤ *p_holm_* < 0.05; **: 0.001 ≤ *p_holm_* < 0.01; ***: *p_holm_* < 0.001.

Insofar, CPC1 spatial power was shown to starkly reorganize across subjects such that a smaller subset of brain regions became more prominent in driving the overall spatiotemporal pattern’s power compared to wakefulness. But are these dominant regions consistently similar across subjects? To address this question, we computed the Jaccard similarity index (**Methods: Similarity in spatial distribution of regional power**) between subjects based on the spatial distribution of their regional powers contributing to CPC1. Intuitively, the higher the similarity index between two individuals is, the larger the overlap between their respective sets of brain regions dominating the wave’s overall power. Under diminished states, there was a significant decrease in inter-subject similarity, particularly under propofol anesthesia. What is more, the similarity index progressively decreased with increasing anesthesia depth (**Figure 6B**; **Supplementary Table 1**). In contrast, no significant changes in similarity were observed during the transition to sleep. These findings indicate that under propofol anesthesia there was an increase in heterogeneity and disagreement across subjects as to which brain regions dominated the overall power of the CPC1 wave.

Overall, these results demonstrate that during diminished states of consciousness, CPC1 spatial power underwent a marked reorganization such that a smaller subset of brain regions became more prominent in driving the overall spatiotemporal pattern’s power compared to wakefulness. The regions that became dominant, however, were highly heterogeneous across subjects, pointing to more fragmented, idiosyncratic, and less stereotyped wave dynamics during diminished states.

#### D. Psychedelic states of consciousness tend to promote uniform spatial distributions of traveling wave power

In contrast to sleep and anesthesia, psychedelic datasets exhibited comparatively varied spatial redistribution effects (**Supplementary Figure 7**). Even though paired within-subject comparisons identified a significant reduction in CPC1’s Gini coefficient under psychedelic states (*t*=-2.1; *p*=0.036), this association did not survive correction for multiple comparisons (*p_holm_*=0.15) (**Figure 7D, E**). Similarly, mixed-effects modeling did not reveal significant pre-post changes in spatial inequality for any CPC (**Figure 7F**). Therefore, although the distribution of regional power contributing to CPC1 tended to become more spatially uniform under psychedelic states, this effect was weaker and occurred in the opposite direction to the increased spatial concentration observed during sleep and anesthesia.

Moreover, inter-subject similarity analyses revealed either no significant changes or increased similarity under psychedelic states, further pointing towards more uniform spatial power distributions. While LSD, DMT, and ketamine showed no significant changes in Jaccard similarity relative to wakefulness, psilocybin and nitrous oxide exhibited significant increases in similarity (**Figure 6B**; **Supplementary Table 1**), suggesting greater consistency across individuals regarding which regions contributed most to CPC1’s overall power. Taken together with the trend towards reduced Gini coefficients, these findings suggest that psychedelic states might generally promote more uniform and less individually differentiated spatial power distributions towards the global wave CPC1.

## DISCUSSION

In this work, we mapped the dominant propagating patterns of BOLD activity across the human brain and examined how their spatial and temporal properties changed across diminished and psychedelic states of consciousness. Using CPCA, we identified four dominant propagation motifs: a global synchronized wave supporting unimodal-transmodal propagation (CPC1), an anti-correlated unimodal-transmodal pattern (CPC2), an anti-correlated task-positive/task-negative pattern (CPC3), and an anti-correlated visual-somatomotor pattern (CPC4). Among these, the global wave CPC1 was markedly reconfigured across both diminished and psychedelic states of consciousness, compared to baseline wakefulness, but in opposite directions: in diminished states, the time it took for the wave to propagate across brain regions—a metric referred to as traveling component duration—increased, whereas in psychedelic states it decreased. Diminished states were also accompanied by a spatial redistribution of CPC1 power, such that a smaller and more variable—across individuals—subset of brain regions became more prominent in driving the overall spatiotemporal pattern’s power compared to wakefulness, indicating the emergence of more fragmented, idiosyncratic, and less stereotyped wave dynamics. In contrast, psychedelic states tended to promote more uniform and less individually differentiated spatial power distributions towards CPC1. Beyond the global wave, diminished states were additionally characterized by increases in traveling component duration along the unimodal-transmodal CPC2 pattern, whereas psychedelic states were characterized by decreases in traveling component duration along the task-positive/task-negative CPC3 pattern, revealing state of consciousness-specific alterations in macroscale propagation dynamics.

The propagation patterns identified in our study closely match those reported in previous studies that applied similar methodologies on resting-state fMRI data in both children^15^ and adults^14,20–22^, reiterating how fMRI dynamics can be parsimoniously captured by a small number of dominant spatiotemporal patterns of propagation. Our work builds on these foundational findings in several ways. Notably, this is to our knowledge the first study to comprehensively compare BOLD-derived propagation patterns of functional activity across both diminished and psychedelic states of consciousness within a unified analytical framework. Building on this comparison, we further advance the literature by systematically characterizing how these propagation patterns reorganize across a broad spectrum of psychedelic states, a topic that has remained largely unexplored. To extend previous characterizations of such patterns which have largely focused on temporal features, we quantified wave properties using two complementary approaches that capture both the temporal dynamics and spatial organization of the propagation patterns. In addition, whereas prior work extracted these spatiotemporal patterns at the population level during wakefulness, we extracted them at the individual level across different states of consciousness, enabling subject- and state-specific characterization of propagation dynamics. This approach allowed us to directly quantify within-subject changes in propagation dynamics, thereby reducing the influence of between-subject variability. To ensure that the individual-derived spatiotemporal patterns were then comparable across different states of consciousness, we aligned them to reference patterns derived from the HCP cohort (**Methods: Extraction of spatiotemporal propagation patterns**; **Supplementary Figure 4**) and compared their spatiotemporal properties.

Critically, our work further emphasizes that although two of these patterns (CPC1 and CPC2) capture unimodal-transmodal propagation patterns, they reflect distinct dynamical regimes: CPC1 represents a largely synchronized global wave where unimodal and transmodal networks co-fluctuate, with the power of the wave primarily being driven by unimodal and attentional networks. In contrast, CPC2 reflects an anti-correlated unimodal-transmodal motif in which unimodal and transmodal networks occupy opposing phases of the propagation cycle, with transmodal networks contributing most strongly to the overall power of the wave. Although the unimodal-transmodal gradient is typically treated in literature as a single dynamical mode of information transfer, our results indicate that it can be decomposed into two distinct propagation motifs that differentially organize neural communication.

In addition to propagating along the unimodal-transmodal hierarchy, CPC1 closely tracks the brain’s global signal and strongly aligns with the average latency structure of spontaneous BOLD fluctuations.^14,23^ Beyond its role in functional connectivity organization, multiple lines of evidence indicate that the global signal is intricately woven into arousal-related physiology and neuromodulatory systems.^24^ For example, a study examining the prevalence of CPC1 relative to other CPCs during prolonged resting-state fMRI scans showed that CPC1 increasingly dominated the signal, with both its prevalence and power rising over the course of the scan, as individuals presumably transitioned to lower arousal and vigilance levels.^25^ In further support of this notion, spontaneous fluctuations in arousal have also been associated with global waves of BOLD activity slowly propagating in parallel throughout the neocortex, thalamus, striatum, and cerebellum.^26,27^ More broadly, recent work demonstrated that the global fMRI signal is tightly coupled to autonomic physiological processes, including fluctuations in respiratory volume, heart rate, and pupil diameter,^28–30^ and reflects coordinated neural activity likely regulated by the basal forebrain:^31^ a group of structures heavily involved in arousal control.^32^

It is therefore not surprising that the duration of the CPC1 propagation component captured in our study was starkly altered across diminished (NREM sleep, propofol anesthesia) and psychedelic states of consciousness where level and content of consciousness are profoundly affected. Importantly, however, the global wave CPC1 was reconfigured in opposite directions across these states: its traveling component duration consistently lengthened in diminished states, while it shortened in psychedelic states. These opposing dynamics could point towards diminished and psychedelic states as potentially capitalizing on the global signal to process conscious content in distinct ways. Case in point, slow cortical potentials—the low-frequency (< 4 Hz) end of field potentials closely related to global BOLD fluctuations—have been proposed to influence whether incoming stimuli reach conscious awareness by modulating higher-frequency activity and fostering large-scale information integration across the brain.^33^ Subsequent work has shown that global waves propagating across the brain might help coordinate activation of large-scale functional networks, synchronizing their activity with fluctuating levels of arousal,^26^ while diminished states of consciousness including sleep and propofol anesthesia are accompanied by loss of global temporal coordination, with distinct patterns of decoupling between local regions and global activity distinguishing between different forms of unconsciousness.^34^ Taken together, these findings suggest that global brain dynamics might provide a temporal scaffold for integrating and updating conscious content. Therefore, the marked prolongation of global signal propagation in diminished states observed in our work could reflect a slowing of this integrative scaffold, effectively lengthening the temporal window over which information is coordinated,^35^ thereby reducing responsiveness to environmental stimuli and attenuating experiential richness. In contrast, the observed accelerated global propagation in psychedelic states may shorten this temporal integration window, promoting more rapid updating of neural representations and a more fluid phenomenology. This interpretation is further supported by recent work identifying accelerated traveling waves under psilocybin, with faster wave propagation being associated with greater psychedelic experience intensity.^36^ Broadly, our study thus posits that the global wave CPC1 could indeed represent a parsimonious dynamical signature with the capacity to differentiate these opposing regimes of consciousness.

To benchmark our approach against more conventional spectral measures of global brain activity, we also computed the mean brain signal across all regions for each state of consciousness, estimated its power spectral density, and assessed shifts in mean frequency between baseline wakefulness and each altered state. These analyses revealed no significant differences in mean frequency across states (**Supplementary Figure 9**). Therefore, traveling component duration might provide greater sensitivity to state of consciousness-dependent changes in BOLD propagation dynamics than traditional frequency-based summaries of the global signal.

Besides these temporal changes, there was also a pronounced spatial redistribution of CPC1 power (squared magnitude of the corresponding spatial vector) across regions during diminished states of consciousness, wherein a smaller subset of brain regions became more relevant in driving the overall spatiotemporal pattern’s power, compared to wakefulness. This finding could reflect the reduction in the dimensionality of neural activity observed during sleep and propofol anesthesia, whereby brain dynamics become increasingly constrained to a limited set of dominant modes.^37–39^ Importantly, similarity analyses revealed that these dominant regions varied substantially across individuals, indicating more fragmented, idiosyncratic, and less stereotyped wave dynamics during diminished states. Consequently, the increased concentration of CPC1 power within a restricted subset of regions, combined with the reduced inter-individual similarity, may reflect the emergence of a lower-dimensional and less-integrated energy landscape, consistent with reductions in global integration^40–45^ and increases in structure-function coupling strength^46–51^ reported during diminished states of consciousness, wherein global activity patterns are driven by fewer effective degrees of freedom. Such reduced integration may, in turn, destabilize canonical propagation architectures, leading to more heterogeneous and less stereotyped wave dynamics across individuals.

In contrast, a trend towards the opposite direction emerged across psychedelic states, where the spatial distribution of CPC1 power tended to become more evenly distributed across brain regions, indicating that a broader set of regions contributed to the global wave’s overall power. What is more, similarity analyses revealed either no change or increased inter-subject similarity under some psychedelic compounds including psilocybin and nitrous oxide. This result could further suggest that regional contributions to CPC1 power appeared to be more uniformly expressed across individuals. Such broader and homogeneous participation of brain regions in the global wave might be indicative of increased large-scale coordination, and therefore reflect a regime of enhanced integration across the brain—consistent with prior literature reporting decreased brain modularity, flattened hierarchical organization, and increased global connectivity and integration under psychedelic compounds such as LSD,^52–54^ DMT,^55^ psilocybin.^54,56^ Additionally, our results complement prior work reporting a relatively preserved perturbational complexity index^57^—a measure of the brain’s capacity to sustain both differentiated and integrated responses to external perturbations^58^—during psychedelic states such as those induced by psilocybin^59^ and ketamine,^60^ in contrast to robust decreases in this metric observed during sleep and anesthesia.^58,61^ Overall, in our work, diminished states of consciousness were characterized by slower CPC1 traveling dynamics and a more spatially concentrated—yet highly heterogeneous across individuals—organization of CPC1 activity, whereas psychedelic states predominantly accelerated CPC1 temporal propagation while appearing to promote a more spatially distributed and integrated pattern of wave propagation across the brain.

Besides altering the spatiotemporal properties of the global wave CPC1 in opposite directions, another notable finding in this work was that each consciousness regime was also characterized by distinct wave profiles: diminished states of consciousness were specifically associated with increased traveling component duration along the unimodal-transmodal CPC2 wave, while psychedelic states were associated with decreased traveling component duration along the task-positive/task-negative CPC3 wave. These observations suggest that altered states of consciousness do not only modulate global brain dynamics, but also selectively reshape propagation along specific well-established hierarchies, depending on the underlying state (diminished versus psychedelic). The unimodal-transmodal hierarchy reflects a prominent gradient of functional organization,^19^ separating sensory and motor regions from higher-order association cortices, and has been proposed to capture fundamental differences in temporal integration and information processing across the brain.^35,62–64^ Within this framework, the slowing of propagation along the CPC2 unimodal-transmodal axis during diminished states may reflect a reduced capacity for hierarchical integration between sensory and transmodal systems, consistent with previous observations that sleep and anesthesia disrupt long-range communication and coordinated transfer of information between these systems.^21,40,65–69^ In contrast, the selective acceleration of propagation along the task-positive/task-negative CPC3 axis under psychedelic states—particularly those induced by serotonergic psychedelics—may reflect increases in functional connectivity and integration between the default mode network and task-positive control systems, such as the frontoparietal network.^70–74^ This aligns with the functional roles of these systems, with the DMN supporting introspection^75–77^ and control systems supporting externally focused attention.^70,78^ Indeed, structured propagation changes along biological gradients persisted even after regressing out the global signal (and thus the global propagation pattern CPC1) from the BOLD signal time-series: diminished states continued to show slowing along the unimodal-transmodal axis, whereas psychedelic states continued to exhibit accelerated dynamics along the task-positive/task-negative axis (**Supplementary Analysis 1**). Together, these findings suggest that diminished and psychedelic states not only modulate global dynamics in opposite directions, but also differentially impact the propagation of neural activity along key gradients of brain organization, offering insight into how these distinct consciousness regimes influence the processing of conscious content.

There are several methodological considerations worth noting. First, we extracted the dominant spatiotemporal patterns of propagation in each individual separately at each state of consciousness, and then identified those that best matched the reference patterns derived from the HCP cohort. Although this process allowed us to directly compare pattern properties at the individual and state level, it assumed that comparable propagation patterns were present across all subjects and states. This assumption might not always be the case—especially across varying drug concentrations or across altered states such as deep sedation that are phenotypically distinct from wakefulness—and might generate propagation patterns that deviate from those derived from the wakeful HCP data. In this study, however, the matching between the altered states and the wakeful references did yield very consistent results (as can be seen from **Supplementary Figure 4**). Another methodological consideration is our definition of traveling component duration. In our approach, the traveling component was inferred indirectly from the spatial phase range of the spatiotemporal pattern, which can be sensitive to phase wrapping and to regions with aberrantly low amplitude where phase estimates become unreliable. To address the former issue, we applied a reference-independent phase span procedure to compute phase ranges (**Methods: Traveling component duration of spatiotemporal patterns**); to mitigate the latter issue, we excluded brain regions with exceedingly low amplitude before estimating the spatial phase range (**Methods: Traveling component duration of spatiotemporal patterns**). Critically, it should also be noted that traveling component duration represents the time (in seconds) needed for the wave to traverse the range of phase offsets observed across brain regions within the spatiotemporal pattern. It should thus be interpreted as a summary index of the extent of propagation within a spatiotemporal pattern, rather than a direct estimate of wave-transit duration. More broadly, the interpretation of what each spatiotemporal pattern/CPC biologically captures warrants careful consideration. Although CPCA identifies dominant spatiotemporal propagation patterns, the physiological and neuromodulatory processes shaping each component remain incompletely understood, and their relative contributions may vary across states of consciousness. For instance, even though the global CPC1 wave appears closely linked to arousal-related global signal fluctuations, the global BOLD signal is also known to capture signals of non-neuronal origin.^29,79–81^ Therefore, disentangling the respective neuromodulatory and physiological contributions to each CPC represents an important direction for future work.

Lastly, our classification of altered states also warrants qualification. In this work, we adopted an operational definition in which nitrous oxide and ketamine are considered non-classical psychedelics, based on their capacity to reliably induce marked alterations in conscious experience that overlap phenomenologically with those produced by classical psychedelics.^74^ We emphasize that this classification is limited and heuristic, as both compounds differ in their primary mechanisms of action—most notably NMDA receptor antagonism^82^—compared to the serotonergic mechanisms that define classical psychedelics.^83^ Accordingly, our usage should not be interpreted as implying pharmacological equivalence, but rather as a pragmatic grouping based on partially shared experiential and dynamical features.

In conclusion, we characterized the dominant spatiotemporal propagation patterns of the BOLD signal and examined how these patterns change across altered states of consciousness. To that end, we analyzed diminished states including NREM sleep and propofol anesthesia, as well as psychedelic states induced by classical psychedelic compounds (LSD, DMT, psilocybin) and non-classical psychedelic compounds (nitrous oxide, ketamine). Across these states, we identified a global brain wave whose propagation duration and spatial organization became starkly reconfigured, distinguishing diminished states from psychedelic states. Furthermore, these states exerted dissociable effects on the brain’s principal axes of functional organization: diminished states principally altered propagation along the unimodal-transmodal axis, whereas psychedelic states appeared to modify propagation along the task-positive/task-negative axis. Collectively, diminished and psychedelic states are characterized not only by changes in global brain dynamics, but also by differential alterations in the propagation of neural activity along key gradients of brain organization.

## METHODS

### DATASETS

#### Human Connectome Project

A sample of 100 unrelated healthy subjects (54% female; age = 29.1 ± 3.7 years; age range = 22 - 36 years) was drawn from the publicly available S1200 release of the Human Connectome Project (HCP).^16^ Subjects were scanned on a customized Siemens Connectome Skyra 3T scanner (32-channel Siemens head coil) and underwent high-resolution 3T MRI, including a T1-weighted (3D Multi-echo Magnetization-Prepared Rapid Gradient Echo [MEMPRAGE] sequence; voxel size: 0.7 mm isotropic; repetition time [TR]: 2400 ms; echo time [TE]: 2.14 ms) and resting-state fMRI (gradient-echo echo-planar imaging [EPI] sequence; voxel size: 2 mm isotropic; TR: 720 ms; TE: 33.1 ms) sequences.^84,85^ Four resting-state fMRI sequences were acquired in two fMRI sessions (rest1 and rest2). Each session comprised of two sequences, each consisting of 1200 volumes and lasting 14 min and 33 sec, acquired using opposite phase encoding directions (left-to-right and right-to-left) to minimize distortion. To mitigate phase encoding-related biases, the two sequences were temporally concatenated, yielding a single fMRI signal time-series per session. For the purposes of this work, we only used the fMRI signal time-series corresponding to the first session (rest1: 2400 volumes; total duration: 29 min and 6 sec). Informed consent was obtained from all subjects, and the procedures were approved by the Washington University Institutional Review Board (IRB).

#### Propofol Dataset 1 (light and intermediate sedation): PPF 1.0 and PPF 1.9

Healthy participants (n = 15; 6 females; age range = 19 – 35 years) were recruited at the Medical College of Wisconsin, where they were sedated with the anesthetic agent propofol, while undergoing MRI, as previously described.^86,87^ The participants first completed a resting-state fMRI scan while awake (baseline). Propofol was then administered using a bolus dose followed by target-controlled continuous infusion to achieve plasma concentrations corresponding to light sedation (0.98 ± 0.18 μg/mL; lethargic response to verbal commands) and intermediate sedation (1.88 ± 0.24 μg/mL; no response to verbal commands or mild physical prodding). Resting-state fMRI scans were acquired at each sedation level, followed by a final recovery scan once behavioral responsiveness had returned. Sedation depth was continuously monitored and infusion rates were manually adjusted to maintain steady-state drug levels. Standard physiological monitoring (electrocardiogram, blood pressure, pulse oximetry, end-tidal carbon dioxide gas) and supplemental oxygen via nasal cannula were provided throughout.

MRI data were collected on a 3T Signa GE 750 scanner (GE Healthcare, Waukesha, WI, USA) with a 32-channel transmit/receive head coil. Each of the four resting-state fMRI scans acquired (wakefulness, light sedation, intermediate sedation, recovery) lasted 15 minutes (TR = 2 s, TE = 25 ms, slice thickness = 3.5 mm, in-plane resolution = 3.5 × 3.5 mm^2^). High-resolution anatomical images (three-dimensional spoiled gradient-recalled echo) were also acquired (TE/TR/TI = 8.2/3.2/450 ms; slice thickness = 1 mm). Informed consent was obtained from all participants, and the procedures were approved by the Medical College of Wisconsin IRB.

#### Propofol Dataset 2 (intermediate sedation): PPF 2.2

Healthy participants (n = 26; 13 females; age range = 19 – 34 years) were recruited at the University of Michigan Hospital, in Ann Arbor, where they were sedated with the anesthetic agent propofol, while undergoing MRI. This publicly available dataset (OpenNeuro accession number ds006623) has been previously described in detail^88^ and utilized in multiple prior studies.^89–92^

The participants first completed a resting-state fMRI scan lasting 10 minutes, while awake and with their eyes closed (Rest 1). Task-based fMRI data were then acquired across four runs: a 15-minute awake baseline scan (Imagery 1), a 30-minute scan during propofol exposure and loss of consciousness (Imagery 2), a 30-minute post-infusion scan where the individuals began regaining consciousness (Imagery 3), and a 15-minute recovery awake baseline scan (Imagery 4). Across these task-based runs, participants performed guided mental imagery and motor response tasks, including tennis imagery, spatial navigation imagery, hand squeezing imagery, and actual hand squeezing tasks. Behavioral responses to action commands were used to delineate pre-loss of responsiveness, loss of responsiveness, and recovery of responsiveness periods. After the task-based fMRIs had been acquired, the participants repeated one more 10-minute long resting-state fMRI scan (Rest 2). This protocol allowed the acquisition of fMRI data spanning wakefulness, induction to unconsciousness, unconsciousness, recovery from unconsciousness, and wakefulness again. Four participants were excluded from subsequent group analyses due to excessive movement in the scanner during fMRI acquisition.

Propofol was administered during Imagery 2 via a target-controlled intravenous bolus followed by a continuous rate infusion. Dosages (bolus and infusion rates) were increased every five minutes in increments of 0.4 μg/mL, until the participant displayed loss of behavioral responsiveness. The final target concentration (2.3 ± 0.5 μg/mL) was maintained for 21.6 ± 10.2 minutes. Afterwards, propofol infusion was stopped to allow spontaneous recovery of consciousness (Imagery 3). Propofol administration was closely monitored by on-site anesthesiologists who continuously monitored vital signs (spontaneous breathing, end-tidal carbon dioxide gas, heart rate, pulse oximetry, electrocardiogram, and arterial pressure); supplemental oxygen was also provided via a nasal cannula.

MRI data were collected on a 3T Philips scanner equipped with a standard 32-channel transmit/receive head coil. Whole-brain functional images were acquired using a gradient-echo echo-planar imaging (EPI) sequence with parameters: 28 slices, TR/TE = 800/25 ms via multiband (MB) acquisition (MB factor = 4), slice thickness = 4 mm, in-plane resolution = 3.4 × 3.4 mm^2^. Notably, six participants were scanned with slightly modified parameters: 21 slices, TR/TE = 800/25 ms, MB factor = 3, and slice thickness = 6 mm, as their data were collected prior to MRI hardware upgrades.^88^ High-resolution anatomical images were also captured (T1-weighted spoiled gradient recalled echo sequence: TR = 8.1 s, TE = 3.7 ms, slice thickness = 1.0 mm with no gap). Informed consent was obtained from all participants, and the procedures were approved by the University of Michigan IRB.

#### Propofol Dataset 3 (deep sedation): PPF 2.7

This next dataset was also acquired at the University of Michigan Hospital, in Ann Arbor. Healthy participants (n = 30; 20 females; age range = 18 – 38 years) were once again recruited to undergo MRI while receiving propofol administration, as previously described.^91^ Each participant completed a series of eight fMRI scans, each lasting 16 minutes, spanning wakefulness, propofol-induced sedation, and recovery. During baseline wakefulness, participants completed a resting-state fMRI scan (Rest 1) and a music-listening fMRI scan (Music 1), followed by a behavioral assessment scan during anesthetic induction. After losing behavioral responsiveness, participants underwent a second set of resting-state (Rest 2) and music-listening (Music 2) fMRI scans. This was succeeded by another behavioral scan captured during the recovery of consciousness phase, after propofol infusion had been terminated. Subsequently, participants performed one final set of resting-state (Rest 3) and music-listening (Music 3) fMRI scans once behavioral responsiveness had returned. Rest 1/Music 1 defined the baseline wakefulness state, Rest 2/Music 2 the loss of responsiveness (i.e., unconsciousness) state, and Rest 3/Music 3 the recovery wakefulness state. Five individuals were excluded from subsequent group analyses due to excessive movement in the scanner (n = 2), truncated field-of-view in fMRI during unconsciousness (n = 1), or incomplete fMRI data collection during unconsciousness (n = 2).

The anesthetic administration and monitoring were similar with that of the previous propofol dataset, with a few exceptions: instead of steadily increasing the anesthetic dosage until the participant displayed loss of behavioral responsiveness, propofol infusion in this experimental protocol was manually adjusted (during the first behavioral assessment scan) to reach specific target effect-site concentrations of 1.5, 2.0, 2.5, and 3.0 μg/mL. To accurately pinpoint the propofol concentration at which each individual lost behavioral responsiveness, each target effect-site concentrations was maintained for 4 minutes followed by manual adjustment of the anesthetic dosage. To mitigate any head motion-related artifacts, the effect-site propofol concentration was kept at one level higher (+0.5 μg/mL) than the concentration associated with loss of responsiveness (average value across all participants: 2.7 ± 0.5 μg/mL), and was maintained for approximately 32 minutes. Past this point, the propofol infusion was stopped during the second behavioral assessment scan to allow spontaneous recovery of consciousness.

MRI data were collected on a 3T Philips MRI scanner equipped with a standard 32-channel transmit/receive head coil. Functional images were acquired using a gradient-echo EPI sequence with parameters: TR/TE = 1400/30 ms via multiband acquisition (MB factor = 4), slice thickness = 2.9 mm, and in-plane resolution = 2.75 × 2.75 mm^2^. High-resolution anatomical images were also captured (T1-weighted spoiled gradient recalled echo sequence: TR = 8.1 s, TE = 3.7 ms, slice thickness = 1.0 mm). Informed consent was obtained from all participants, and the procedures were approved by the University of Michigan IRB.

#### Natural Sleep

Healthy participants (n = 33; 16 females; age = 22.1 ± 3.2 years) were recruited at Pennsylvania State University to investigate spontaneous brain activity across distinct sleep stages, using simultaneous EEG-fMRI acquisition. Data were acquired across wakefulness and three progressively deeper NREM sleep stages (N1, N2, and N3); sleep stages were identified via EEG by a registered polysomnographic technologist, as previously described.^93,94^ Given that only three participants entered N3 sleep during this acquisition, we only considered data from N1 (light sleep) and N2 (deeper sleep) stages in our analyses. This dataset has been made publicly available through OpenNeuro (accession number ds003768).

MRI data were collected on a 3T Prisma Siemens Fit scanner with a Siemens 20-channel receive-array coil. Functional MRI data were acquired using an EPI sequence with parameters: TR = 2100 msec, TE = 25 msec, slice thickness = 4 mm, in-plane resolution = 3 x 3 mm^2^. Structural MRI data were also acquired using a magnetization-prepared rapid acquisition gradient echo (MPRAGE) sequence (TR/TE/TI = 2300/2.28/900 msec, voxel size = 1 x 1 x 1 mm^3^). Informed consent was obtained from all participants, and the procedures were approved by the Pennsylvania State University IRB.

#### LSD

In this publicly available dataset (OpenNeuro accession number ds003059),^95,96^ healthy participants (n = 15; 4 females, age = 30.5 ± 8 years) attended two scanning sessions (LSD and placebo) at least two weeks apart in a balanced, within-subjects design. Each participant received bolus injections of LSD (75 μg in 10 mL saline) or placebo (10 mL saline) over two minutes, while they were instructed to close their eyes and relax. Following an acclimatization period of 60 minutes, three fMRI scans were acquired: an eyes-closed resting state, a resting state scan while the individuals listened to music, and another eyes-closed resting state scan. All three placebo fMRI scans were used to define a baseline state, while all three LSD fMRI scans were used to define the LSD state.

MRI data were collected using a 3T GE HDx scanner. Functional scans were acquired using a gradient echo planar imaging sequence, with parameters: TR/TE = 2000/35 msec, voxel size: 3.4 x 3.4 x 3.4 mm^3^). Anatomical scans were also acquired using a 3D fast spoiled gradient echo sequence (TR/TE = 7.9/3 msec). Informed consent was obtained from all participants; this study was approved by the National Research Ethics Service committee London – West London, and was conducted in accordance with the revised declaration of Helsinki (2000), the International Committee on Harmonization Good Clinical Practice guidelines, and the National Health Service Research Governance Framework.

#### DMT

Healthy participants (n = 20, 7 females, age = 33.5 ± 7.9 years) were recruited to undergo combined fMRI-EEG, in a single-blind, placebo-controlled, counter-balanced design aimed to examine the acute effects of DMT administration in the brain, as previously described.^55^

Specifically, volunteers completed two sessions at the Imperial College Clinical Imaging Facility, spaced two weeks apart. During the first (task-free) session, participants received intravenous administration of either placebo (10 mL sterile saline) or DMT (20 mg), delivered over 30 seconds and followed by a 10 mL saline flush over 15 seconds, in a counter-balanced order. This session consisted of a continuous resting-state fMRI scan lasting 28 minutes, with DMT/placebo administered at the end of the 8^th^ minute and scanning continuing for 20 minutes post-injection. The second session followed the same procedure with the only difference that participants were audio-prompted to provide minute-by-minute verbal ratings of their subjective drug intensity while in the scanner. In this work, we report results pertaining to the resting-state scans obtained from the first task-free session, as in previous work.^55^ Specifically, the first 8 minutes of the acquired scan were used to define baseline, while the 8 minutes post-DMT administration were used to define the DMT state.

MRI data were collected using a 3T Siemens Magnetom Verio Syngo MR B17 scanner equipped with a 12-channel head coil. Functional imaging was performed using a T2*-weighted gradient EPI sequence with parameters: TR = 2000 msec, TE = 30 msec, duration = 28.06 min, voxel size = 3.0 × 3.0 × 3.0 mm^3^, and interslice distance = 0 mm. This study was approved by the National Research Ethics Service committee London – Brent and the Health Research Authority, and was conducted under the guidelines of the revised Declaration of Helsinki (2000), the International Committee on Harmonization Good Clinical Practices guidelines, and the National Health Service Research Governance Framework.

#### Psilocybin

Healthy participants (n = 7, age range = 18 – 45 years) were recruited at Washington University in Saint Louis in a study poised to evaluate differences in functional connectivity before, during, and after psilocybin administration. More details regarding this dataset and how to access it can be found in the corresponding manuscript.^97^ The study employed a randomized cross-over design wherein participants underwent MRI during non-drug sessions, as well as drug sessions where they were administered either high-dose psilocybin (25 mg) or an active placebo control (40 mg methylphenidate). For the purposes of this study, we only focused on the baseline non-drug and psilocybin sessions.

MRI data were collected using a 3T Siemens Prisma scanner. Resting state fMRI scans were acquired using a multi-echo EPI sequence (TR = 1761 msec, TEs = 14.2, 38.93, 63.66, 88.39, 113.12 msec, MB factor = 6, voxel size: 2 x 2 x 2 mm^3^, duration = 15 min:49 sec). To ensure that results obtained from this dataset were comparable to those of other datasets, we only utilized the sequences corresponding to TE = 38.93 msec. High resolution anatomical (T1- and T2-weighted) scans were also collected (voxel size: 0.9 x 0.9 x 0.9 mm^3^). Informed consent was obtained from all participants, and the procedures were approved by the Washington University in Saint Louis IRB.

#### Ketamine

Healthy participants (n = 12, 5 females, age = 41.4 ± 8.6 years) were enrolled at Huashan Hospital, Fudan University, to examine the effects of ketamine on brain function, as previously described.^74,98^ Participants first underwent a 10-minute long baseline resting-state fMRI (with the exception of 2 individuals for whom the scan lasted 6 and 11 min, respectively). They then received an infusion of ketamine (0.5 mg/kg per minute) for 10 minutes and another infusion of ketamine (0.1 mg/kg per minute) for another 10 minutes, except for two participants who received only the second infusion. The ketamine infusion was then discontinued, allowing the participants to naturally regain consciousness. This experimental protocol essentially consisted of two phases: (1) a subanesthetic ketamine administration period typically associated with psychedelic experiences, and (2) a period associated with ketamine-induced loss of behavioral responsiveness. Similar to previous work,^74^ we only focused on the subanesthetic ketamine administration period.

MRI data were collected using a 3T Siemens MAGNETOM scanner equipped with an eight-channel head coil. Whole-brain functional images were acquired using a gradient-echo EPI sequence with parameters: TR/TE = 2000/30 msec, slice thickness = 5 mm. Informed consent was obtained from all participants, and the procedures were approved by the Fudan University’s Huashan Hospital IRB.

#### Nitrous Oxide

Healthy participants (n = 16, 8 females, age = 24.6 ± 3.7 years) were recruited to participate in a study at the University of Michigan Hospital, in Ann Arbor, examining the effects of subanesthetic dosage of nitrous oxide on functional connectivity, as previously reported in detail.^74,99^ Briefly, participants performed a resting-state fMRI scan, followed by an fMRI scan in the presence of visual stimuli, and an fMRI scan while experiencing pressure stimuli via a cuff. They repeated this set of scans before and after exposure to subanesthetic doses of inhaled nitrous oxide (35%). The three scans acquired before nitrous oxide administration were used to define the baseline state, while the same scans acquired after nitrous oxide administration were used to define the altered state. Two individuals were excluded from subsequent group analyses due to excessive movement during fMRI acquisition, as in previous work.^74,99^

MRI data were collected using a 3T Philips Achieva scanner. Each functional image was acquired using a T2*-weighted echo-planar sequence with parameters: TR/TE = 2000/30 msec, and slice thickness = 3 mm. Informed consent was obtained from all participants, and the procedures were approved by the University of Michigan IRB.

### PROCESSING PIPELINES

#### fMRI Processing

The fMRI scans extracted from all aforementioned datasets were analyzed using a previously established pipeline,^89–91,100^ implemented in AFNI (Analysis of Functional NeuroImages; https://afni.nimh.nih.gov). Preprocessing of the fMRI scans included (i) the removal of the first two frames from each fMRI, (ii) slice timing correction to adjust for the temporal differences in slice acquisition, (iii) identification of frames with framewise displacement values greater than 0.4 mm, indicating excessive head motion, (iv) co-registration of the fMRIs to T1-weighted anatomical images, (v) spatial normalization into standardized Talairach stereotactic space (3 x 3 x 3 mm^3^), and (vi) bandpass filtering the fMRI data to 0.01 – 0.1 Hz. Confounding signals were then removed from the preprocessed fMRI data using linear regression. Such signals included time-series of head motion (i.e., framewise displacement) and its temporal derivative, binary framewise displacement time-series indicating which frames were flagged as being of excessive head motion, time-series corresponding to the mean white matter and cerebrospinal fluid signals, as well as linear and non-linear drifts. Frames with excessive motion were replaced with linearly interpolated frames derived from the non-censored neighboring frames, to ensure temporal continuity.

To examine how the inclusion of the global signal in the fMRI data impacted its spatiotemporal propagation patterns, we generated two versions of the fMRI signal: one that included the global signal (without global signal regression [GSR] approach), and one where the global signal and its temporal derivative were regressed out (GSR approach). **Figures 1, 2, 4-7**, **Supplementary Figures 1-9**, and **Supplementary Table 1** show results without GSR, while **Supplementary Figures 10-12** show results with GSR. Results pertaining to the analyses with GSR can be found in **Supplementary Analysis 1**. Denoising of the fMRI signals was subsequently followed by (vii) spatial smoothing with 6 mm full-width-at-half-maximum isotropic Gaussian kernel, and (viii) temporal normalization of the signal to zero mean and unit variance. For both with and without GSR approaches, average fMRI signals were extracted from 400 cortical regions and 50 subcortical regions, derived from previously established atlases,^17,101^ for each individual in each dataset.

For some of our figures (**Figures 1, 2** and **Supplementary Figures 2, 10**), we also display the average fMRI signal for (i) each of seven canonical resting-state functional cortical networks (visual, somatomotor, dorsal attention, ventral attention, limbic, frontoparietal, and default mode) spanning the unimodal-transmodal hierarchy (assignment of each of the 400 cortical regions into its corresponding resting-state system was provided with the cortical atlas),^17^ as well as (ii) several subcortical networks including the hippocampus, amygdala, basal ganglia, and thalamus (assignment of each of the 50 subcortical regions into its corresponding system was provided with the subcortical atlas).^101^

#### Complex Principal Component Analysis

To extract each individual’s spatiotemporal propagation patterns from their fMRI signal at different states of consciousness, we applied a methodology referred to as complex principal component analysis (CPCA).^14,15,102,103^

Recent work has shown that fMRI signals exhibit structured spatiotemporal patterns of propagation (also referred to as waves),^14,15^ and that these patterns can be mathematically extracted using CPCA. Conceptually, CPCA extends conventional PCA into the complex space by transforming the fMRI time-series into analytic signals which encode the signal’s instantaneous amplitude and phase. This representation preserves both the amplitude of signal fluctuations across different brain regions, and their relative temporal shifts as the signal propagates over time. CPCA then decomposes the analytic signal into a series of spatiotemporal modes, each characterized by a spatial pattern and its associated temporal dynamics, enabling the identification of recurring propagation patterns that dynamically unfold across the brain over space and time.

First, an individual’s fMRI time-series matrix *X* ∈ ℝ^*T*×*R*^ (*T*: number of time-points, *R*: number of brain regions) at a given state is Hilbert-transformed. Mathematically, the Hilbert transform *H* converts a real-valued signal *X* into its complex analytic form *X*_*A*_ ∈ ℂ^*T*×*R*^, which encodes information about the signal’s instantaneous amplitude and phase across time (**Equation 1**). Next, the complex fMRI signal is decomposed into its temporal (*U*) and spatial (*V*) constituent matrices via singular value decomposition (**Equation 2**). Under this formulation, *U* ∈ ℂ^*T*×*k*^ and *V* ∈ ℂ^*R*×*k*^ contain the left and right singular vectors, respectively, with *k* = *rank*(*X*_*A*_) ≤ *min*(*T*, *R*), while *Σ* ∈ ℝ^*k*×*k*^ is a diagonal matrix whose entries contain the non-negative singular values ordered in descending magnitude. Conceptually, the columns in *U* represent orthonormal temporal modes, while the columns in *V* represent orthonormal spatial modes; the diagonal entries in *Σ* quantify each mode’s overall contribution. Together, these singular modes constitute orthogonal spatiotemporal patterns, representing complex principal components, and capturing the dominant variance and phase structure in the analytic fMRI signal.

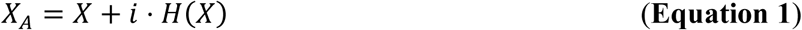

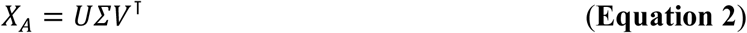

#### Extraction of spatiotemporal propagation patterns

The above formulation enables the extraction of each spatiotemporal pattern’s (i.e., complex principal component’s) temporal and spatial expression. Because our goal was to compare propagation patterns across multiple datasets, we sought to define a common set of reference patterns. To this end, we leveraged the HCP dataset as reference.

Each HCP subject’s fMRI signal time-series (2400 time-points x 450 brain regions) was first Hilbert-transformed to extract the respective complex analytic signal. All subjects’ analytic signals were then concatenated across time (240000 time-points x 450 brain regions), and the resulting matrix was decomposed using CPCA, as described above. This group-level decomposition allowed us to extract the temporal (*U*) and spatial (*V*) vectors corresponding to each complex principal component of the HCP fMRI signal. These vectors contain complex values of the form *α* + *i* ⋅ *β*, with *α*, *β* ∈ ℝ, which can be reformulated using Euler’s formula *e*^*iφ*^ = *cos*(*φ*) + *i* ⋅ *sin*(*φ*) to reveal the signal’s amplitude 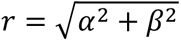 and relative phase angle *φ* = *atan*2(*β*, *α*) expressed in radians *φ* ∈ (−*π*, *π*]. To construct each principal component’s spatiotemporal propagation pattern during a full cycle, we adopted a previously validated approach:^15^ we first divided the full cycle ranging from −*π* to *π* radians into 32 equally spaced phase bins, each spanning 2*π*⁄32 radians. For each complex principal component, time points were assigned to these bins based on the phase of their complex temporal values. Within each phase bin, we averaged the complex values in the corresponding column of *U*, scaled by the component’s singular value in *Σ*, yielding a phase-binned temporal vector of length 32, for each complex principal component. The spatiotemporal propagation pattern of each complex principal component (32 phase bins x 450 brain regions) was then defined as the outer product between its phase-binned temporal vector and its corresponding spatial vector (i.e., column in *V* of length: 450 brain regions). Snippets of the first four complex principal components’ spatiotemporal propagation patterns at different phase bins through the oscillatory cycle are shown in **Figure 1**.

In this work, we focused on the first four complex principal components extracted from the HCP time-series as our reference propagation patterns (henceforth referred to as CPC1-CPC4), as they collectively captured approximately half (48%) of the overall variance of the data. More details regarding the spatiotemporal propagation patterns of CPC1-CPC4 are illustrated in **Figures 1** and **2** and described in the **Results** section.

We next analyzed fMRI data from datasets capturing altered states of consciousness (LSD, DMT, psilocybin, ketamine, nitrous oxide, sleep, and propofol) by applying the CPCA framework to the time-series of each subject, separately for each state and dataset. In many datasets, however, the number of fMRI time-points collected during wakefulness differed from the number of time-points collected during the altered state. To ensure that our results were not biased by differences in the number of time-points across states, we performed a length-matching procedure prior to CPCA decomposition, within each dataset. Specifically, for each state, we computed the median number of time-points across all subjects, and defined the retained length as the minimum of these median values across states. For instance, in a dataset with 10 subjects and two states (wakefulness and altered state), if the median duration was 300 time-points for wakefulness and 500 for the altered state, both states were truncated to 300 time-points. Any excess samples were removed symmetrically from the beginning and end of the time-series to preserve the central temporal segment. We provide below a table summarizing the retained number of subjects and fMRI time-points that were analyzed using our CPCA pipeline, for each dataset:

| **Dataset:** | **Subjects:** | **fMRI time-points:** | **fMRI duration (min):** |
| --- | --- | --- | --- |
| **Propofol Dataset 1** | 14 | 448 | 14.9 |
| **Propofol Dataset 2** | 21 | 770 | 10.3 |
| **Propofol Dataset 3** | 25 | 690 | 16.1 |
| **Sleep** | 12 | 300 | 10.5 |
| **LSD** | 15 | 217 | 7.2 |
| **DMT** | 16 | 240 | 8 |
| **Psilocybin** | 7 | 510 | 15.1 |
| **Ketamine** | 9 | 300 | 10 |
| **Nitrous Oxide** | 14 | 100 | 3.3 |

Applying the CPCA framework on each subject and state produced subject-specific complex principal components and their corresponding spatiotemporal propagation patterns (32 phase bins x 450 brain regions matrix per component). To place these subject-level patterns within a common reference space, we projected them onto the propagation patterns derived from the HCP cohort. For each subject/state, phase-binned spatiotemporal maps were reconstructed as described above and their similarity to the HCP patterns was quantified using an optimal circular correlation procedure. Specifically, for every pair of subject/state and reference components, the subject/state’s phase bin dimension was circularly shifted and the Pearson’s correlation between the flattened spatiotemporal matrices was computed, retaining the maximal correlation and its associated phase shift. We then determined which of the subject/state’s dominant complex principal components (from the first ten) best statistically matched each of the first four HCP components CPC1-CPC4 based on maximal correlation, enforcing unique assignments. The selected subject/state components were phase-aligned to their HCP reference by applying their optimal circular shift, producing aligned spatiotemporal propagation patterns for all downstream analyses.

#### Analysis of spatiotemporal propagation patterns

Following alignment of subject-level spatiotemporal propagation patterns during each state of consciousness to the HCP reference, we next compared their temporal and spatial properties across states. Two principal features were examined: (i) the traveling component duration of each spatiotemporal pattern, that is the portion of the pattern’s overall duration attributed to spatial phase propagation, and (ii) how concentrated each spatiotemporal pattern’s spatial power was across brain regions.

##### Spatiotemporal patterns’ overall duration

Each spatiotemporal pattern’s overall cycle duration was estimated from its corresponding complex principal component’s time-series (i.e., temporal vector in *U*). First, we extracted the instantaneous phase from the complex temporal vector, and temporally unwrapped it to yield a continuous phase trajectory; a full propagation cycle was defined every time an integer multiple of 2*π* was crossed (**Figure 3B**). Since crossings are not always expected to occur exactly at the sampled time-points, we used linear interpolation to estimate crossing times between adjacent samples. This allowed us to optimally deduce duration times in radians with sub-TR precision. Cycle durations were then computed as the temporal differences between consecutive crossings, and the median of these values was designated as the overall cycle duration of the spatiotemporal pattern. Since this number was in radians, we converted it to seconds by multiplying it with that specific dataset’s TR.

##### Traveling component duration of spatiotemporal patterns

To isolate the traveling component of each spatiotemporal pattern, we next estimated the range of spatial phase values derived from the corresponding complex principal component’s spatial vector (i.e., column in *V*). First, we extracted the phase and amplitude of each brain region. Because phase estimates become unreliable when amplitudes are very small,^104^ regions with exceedingly low amplitudes were excluded from this calculation. To that end, the median amplitude across regions was computed and the spread of the distribution was estimated using the scaled median absolute deviation (*SciPy*’s median_abs_deviation tool); regions whose amplitudes fell more than three robust standard deviations below the median were not considered. Using the remaining regions, the spatial phase span was estimated in a reference-independent manner: each region was, in turn, treated as a reference, and the phases of all regions were expressed relative to that reference and wrapped to the interval (−*π*, *π*]. For each reference choice, the difference between the maximum and minimum phase was computed, yielding a phase span. The median of all computed phase spans across all reference choices was defined as the spatial phase range, and this quantity was divided by 2*π* to express it as a fraction of a full cycle (**Figure 3C**). Subsequently, the traveling component duration of the spatiotemporal pattern was defined as this spatial phase range fraction multiplied by the spatiotemporal pattern’s overall duration (**Figure 3D**), reflecting the portion of the cycle that was spent propagating across space, in seconds.

##### Spatial distribution of regional power

We next computed the spatial distribution of regional contributions to each complex principal component. For each subject, state of consciousness, and aligned complex principal component, each brain region’s power was computed as the squared magnitude of the corresponding spatial vector (i.e., |*V*|^2^), reflecting the energy contribution of each brain region to the component (i.e., spatiotemporal pattern), independent of phase. Power values were then normalized by the total power across all brain regions within each component.

To quantify the spatial concentration of regional contributions within each complex principal component, we computed the Gini coefficient (*G*) of the normalized regional power distribution (**Equation 3**; **Figure 3E**). In essence, the Gini coefficient is computed by first rank-ordering the regional power values *x*_*i*_ in ascending order (where *i* indexes brain regions and *n* denotes the total number of regions; here, *n* = 450) and then measuring the extent to which cumulative power deviates from a perfectly uniform distribution across regions.

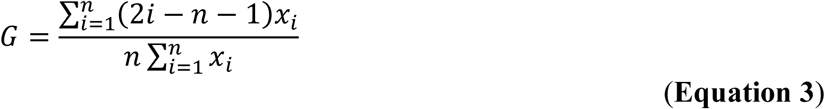

This metric ranges from 0 to 1, with lower values indicating that power is evenly distributed across brain regions and higher values reflecting greater concentration of power within a smaller subset of regions. Intuitively, the Gini coefficient measures how regionally focal versus distributed the spatial power contributions are within each spatiotemporal propagation pattern.

##### Similarity in spatial distribution of regional power

To further quantify the consistency of dominant brain regions (i.e., brain regions with the top X % of regional power within a wave, as defined in the previous section), we computed the Jaccard similarity index between subjects based on the spatial distribution of their regional power for each complex principal component. For each subject and each state of consciousness, we defined a binary mask across brain regions, where regions with the top X % of regional power within a wave (here X = 50) were labeled as 1 and all the rest as 0. Pairwise Jaccard similarity was subsequently computed between each pair of subjects within a state, using the following formula:

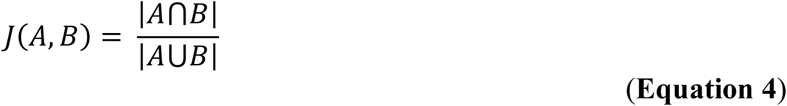

Here, *A* and *B* denote the binary masks across regions for any two subjects, with the numerator and denominator denoting the size of their intersection and union, respectively. Therefore, a Jaccard similarity index of 0 indicates that there is no overlap in dominant regions between two subjects, while a similarity index of 1 indicates that the sets of dominant regions were identical across these two subjects. We computed the pairwise similarity indices across all subject pairs within each state of consciousness, and the resulting distributions were compared between wakefulness and altered states using paired *t*-tests, corrected for multiple comparisons (Benjamini-Hochberg false-discovery rate).

### STATISTICAL ANALYSES

#### Correlation between spatiotemporal patterns and functional gradients

Spatial correspondence between the HCP-derived spatiotemporal patterns (CPC1-CPC4) and canonical functional gradients was quantified using a bin-wise correlation approach. Functional gradient maps were downloaded from the *neuromaps* library (https://netneurolab.github.io/neuromaps/installation.html).^105^ The spatiotemporal propagation pattern of each complex principal component (32 phase bins x 450 brain regions; the derivation of these maps were described above in section **Extraction of spatiotemporal propagation patterns**) and each functional gradient map were first z-scored across brain regions. To account for the temporal evolution of each spatiotemporal propagation pattern across the 32 phase bins spanning each cycle, we computed the Pearson’s correlation between the map corresponding to each bin (1 x 450 brain regions) and each gradient map. We then identified the phase bin that yielded that maximum correlation, and recorded both the bin ID and corresponding correlation coefficient. Both metrics are reported in **Results: Dominant patterns of spatiotemporal propagation**, as well as **Supplementary Figure 3**.

#### Effect sizes

To quantify the magnitude of state-related differences in traveling component duration and Gini coefficients, we computed paired-sample effect sizes using Hedges’ *g* statistic, implemented by Python’s *pingouin* package. For each dataset and complex principal component, effect sizes were calculated from paired observations (i.e., wakefulness vs. altered-state values). The paired Hedges’s *g* was computed using the standardized mean of the within-subject difference. In contrast to other effect size estimates such as Cohen’s *d*, Hedge’s *g* also corrects for biased estimates of effect size, particularly within small sample sizes.^106^ To estimate uncertainty, we additionally derived non-parametric bootstrap confidence intervals by resampling subject pairs with replacement and recomputing Hedges’ *g* across 5,000 iterations. When reporting effect size analyses in the **Results**, we provide, for each dataset, the paired *t*-test *p*-value corrected for multiple comparisons across all datasets (Benjamini-Hochberg False Discovery Rate; *p_FDR_*), together with the corresponding Hedge’s *g* estimate and its 95% confidence interval. FDR correction for multiple comparisons was specifically applied when evaluating effect-size comparisons across heterogeneous experimental samples (i.e., the 11 psychedelic and diminished-state datasets) to balance sensitivity with Type I error control.

#### Mixed-Effects Models

To assess state-related changes while accounting for the hierarchical structure across our altered states of consciousness datasets, we employed linear mixed-effects modeling implemented in Python using the *statsmodels* library. The dependent variable was defined as the metric of interest (e.g., traveling component duration, overall cycle duration, spatial phase range, or Gini coefficient); fixed effects in the model included state (binary: wakefulness or altered state), complex principal component (categorical: CPC1-CPC4), as well as their interaction. Random intercepts were specified for each dataset to account for baseline differences across samples; random slopes were also specified for each dataset, to capture dataset-specific variability in state-related changes over time. To avoid overfitting given the relatively small and uneven number of subjects across datasets, subject-level random effects (intercepts and slopes) were not included in the mixed-effects model. Formally, the model was specified as:

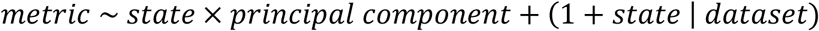

Model-estimated changes are reported in the **Results** section as fixed-effect slope coefficients (*β*), reflecting the estimated difference from wakefulness to altered state of consciousness for the metric of interest, accompanied by 95% confidence intervals, and *p*-values corrected for multiple comparisons (Holm; *p_holm_*) across components. Holm correction was chosen in this case to control for the family-wise error rate across a small set of hypothesis-driven comparisons (i.e., the four complex principal components).

It should be noted that since two of the diminished-states-of-consciousness datasets (Sleep and Propofol Dataset 1) each contained multiple diminished states (Sleep: wakefulness, N1, N2; Propofol Dataset 1: wakefulness, light sedation, intermediate sedation), we included only one diminished condition from each dataset (Sleep: N2, Propofol Dataset 1: intermediate sedation) in the mixed-effects model. This approach avoided repeated sampling from the same participant groups, ensuring that all observations included in the model were independent at the dataset level.

## ACKNOWLEDGEMENTS

This work was supported by NIH grants T32EB035504 (P.F.) and R01GM103894 (A.G.H., Z.H.).

